# Zero-Shot Protein–Ligand Binding Site Prediction from Protein Sequence and SMILES

**DOI:** 10.1101/2025.09.28.679103

**Authors:** Mahdi Pourmirzaei, Salhuldin Alqarghuli, Kai Chen, Mohammadreza Pourmirzaei, Dong Xu

## Abstract

Accurate identification of protein–ligand binding sites is critical for mechanistic biology and drug discovery, yet predictive performance varies substantially across ligand families and data-availability regimes. We present a systematic ligand-aware benchmark and evaluation framework for residue-level binding-site prediction across three settings: overrepresented ligands with abundant examples, underrepresented ligands with limited examples, and zero-shot ligands unseen during training. Using a BioLiP2-derived dataset with ligand-specific sequence clustering, we evaluate a progressive three-stage sequence-based modeling suite designed to study these regimes. Stage 1 trains ligand-specific predictors using a pretrained protein language model (PLM). Stage 2 introduces ligand conditioning through learned embeddings, enabling a single multi-ligand model. Stage 3 replaces these embeddings with representations from a pretrained chemical language model (CLM) operating on SMILES, enabling prediction for previously unseen ligands. In the original three-stage comparison, Stage 2 achieved a higher ligand-macro *F*_1_ than Stage 1 on the overrepresented test set (0.5832 versus 0.4769), although the comparison simultaneously changes predictor architecture, ligand conditioning, and the training regime. Stage 2 also outperformed several sequence- and structure-based baselines. Stage 3 achieves a pooled residue-level *F*_1_ of 0.3109 and a ligand-macro *F*_1_ of 0.2338 on previously unseen ligands, while remaining competitive on represented ligands. Control experiments using constant, mismatched, and scrambled SMILES inputs reduce performance across all three test regimes, providing evidence that the model uses information associated with the supplied ligand input rather than relying solely on generic protein binding-site propensity. Ablations across five PLM scales and multiple CLMs further show that increasing PLM scale generally improves performance, although the trend is not strictly monotonic, whereas CLM scaling produces smaller and less consistent gains. Overall, this study provides a systematic benchmark for quantifying the effect of ligand data availability on residue-level binding-site prediction and for evaluating generalization to previously unseen ligands.

## Introduction

Reliable identification of protein–ligand binding sites plays a critical role in understanding fundamental biological processes such as gene expression regulation, signal transduction pathways, and antigen–antibody interactions Zhao et al. [2020], Dhakal et al. [2022], Xia et al. [2024]. Additionally, precise determination of protein– ligand interaction sites is fundamental for effective drug discovery and rational therapeutic development Xia et al. [2020]. However, pinpointing ligand-binding sites accurately remains challenging, particularly when high-resolution structural data for proteins are unavailable Yu et al. [2013]. Although experimental techniques such as X-ray crystallography, NMR spectroscopy, and cryo-electron microscopy provide structural data for characterizing protein–ligand binding sites, they can be expensive and labor-intensive, underscoring the critical need for computational approaches capable of rapid, high-throughput binding-site prediction Yuan et al. [2022].

Despite notable advancements in computational prediction methodologies for protein–ligand interactions, several limitations persist. Many current computational tools lack versatility, restricting their effectiveness to specific ligand categories Zhao et al. [2020], Xia et al. [2022], Fang et al. [2023], Wang et al. [2024]. This constraint is particularly challenging given the substantial chemical diversity of biologically relevant ligands. Furthermore, data availability varies considerably across ligand types: while some ligands are associated with abundant experimentally characterized protein interactions, many have only limited examples, and others may be entirely absent from the training data. Although recent methods have advanced residue-level binding-site prediction, systematic evaluation of how predictive performance changes across these different ligand data-availability regimes remains limited. A ligand-aware evaluation framework is therefore needed to distinguish performance on data-rich ligands from generalization to data-scarce and previously unseen ligands.

To address this gap, we establish a systematic ligand-aware benchmark and evaluation framework for residue-level protein–ligand binding-site prediction across three distinct regimes: *overrepresented, underrepresented* (few-shot), and *zero-shot* ligands. Protein sequences were clustered within ligand-specific datasets using CD-HIT at 40% sequence identity to reduce sequence redundancy during split construction. We evaluate these regimes using a progressive three-stage sequence-based modeling suite. Stage 1 provides ligand-specific baseline predictors based on a pretrained protein language model (PLM). Stage 2 incorporates learned ligand embeddings to support multiple ligand types within a single model. Stage 3 replaces the learned ligand embeddings with representations derived from SMI-LES using a pretrained chemical language model (CLM), enabling prediction for ligands not observed during training. This framework enables systematic characterization of how residue-level binding-site prediction changes as ligand supervision decreases from data-rich to data-scarce and unseen-ligand settings.

We evaluated the zero-shot model on the modified Stage 3 datasets using SMILES-based ligand inputs. The benchmark includes atomic ions, other inorganic species, and organic or carbon-containing molecules, including larger and organometallic ligands. This evaluated scope should not be interpreted as support for every ligand type in BioLiP2. In the released MolFormer reference evaluation, chemical inputs are limited to 128 tokens, including special tokens, and longer inputs are truncated. This input limit describes the evaluated MolFormer configuration and is not a general limit for all chemical encoders. The evaluated data do not support a universal 100-heavy-atom cutoff. Current SMILES validity, input truncation, and historical encoder compatibility are considered separately.

The main contributions of this work are summarized as follows:

1. **A ligand-aware, three-regime benchmark with withinligand sequence clustering**. We establish a standardized ligand-aware evaluation framework with splits for *overrepresented, underrepresented*, and *zero-shot* ligands, enabling systematic assessment of residue-level binding-site prediction across different levels of ligand data availability.
2. **Systematic evaluation of ligand-aware residue-level prediction**. We use a progressive three-stage modeling suite— comprising ligand-specific predictors, a multi-ligand model conditioned on learned ligand embeddings, and a model conditioned on SMILES-derived CLM representations—to characterize performance as ligand supervision decreases from data-rich to data-scarce and previously unseen ligands.
3. **Controlled analysis of protein and chemical representations**. Through ablations across multiple PLM and CLM configurations and ligand-input controls using constant, mismatched, and scrambled SMILES, we systematically assess the contributions of protein and ligand representations across the three evaluation regimes.

## Related Work

State-of-the-art sequence-based predictors of protein–ligand binding sites increasingly center on pretrained protein language models (PLMs). PLMs address two long-standing challenges: (i) modeling long-range dependencies via transformers and (ii) improving sample efficiency through large-scale pretraining. We group recent PLM-driven approaches by how they adapt pretrained representations to binding-site prediction.

A common line of work keeps the PLM frozen (or lightly tuned) and adds lightweight classifiers. BindEmbed21 Littmann et al. [2021] feeds ProtT5 embeddings into a shallow CNN and combines them with homology-based inference, using class-weighted losses to counter label imbalance. CLAPE-SMB Wang et al. [2024] freezes ESM-2 and employs a five-layer MLP trained with class-balanced focal loss plus a triplet-center term to better separate binding from non-binding sites. For metal ions, LMetalSite Yuan et al. [2022] regularizes PLM embeddings with Gaussian noise, shares transformer layers across tasks, and uses ion-specific MLP heads within a multi-task framework to capture cross-ion commonalities while remaining ligand-aware.

IonPred Essien et al. [2023] instead adopts a generator– discriminator setup, using a masked language model (MLM)-style generator with an ELECTRA-like discriminator to learn residue-substitution regularities, thereby improving data efficiency by reducing reliance on labeled data during fine-tuning.

We developed Prot2Token Pourmirzaei et al. [2025a] using a sequence-only design in which a pretrained ESM-2 encoder provides context to a causal decoder via cross-attention, framing binding-site prediction as next-token generation over sorted residue indices. Each ligand type is represented by a learned *task token* given to the decoder, and the model is trained multi-task over 41 ligand classes. Because conditioning is performed over a closed vocabulary, the approach is effective for a fixed set of ligands but does not directly generalize *zero-shot* to unseen ligands without introducing new tokens and retraining.

LaMPSite Zhang and Xie [2023] augments ESM-2 sequence embeddings with an estimated contact map and pairs them with a graph neural network (GNN) over the ligand’s 2D molecular graph, together with a conformer-derived distance map. A residue–atom interaction module integrates protein contacts and ligand information to refine residue-level binding likelihoods, yielding a sequence-anchored yet ligand-aware predictor.

### Drug–target interaction prediction and unseen-ligand generalization

Related ideas for jointly representing proteins and small molecules have been extensively explored in drug–target interaction (DTI) and binding-affinity prediction. ChemBoost Özçelik et al. [2021] introduced a chemical-language-based approach in which SMILES-derived chemical word representations are combined with protein information for protein–ligand affinity prediction. DeepPurpose Huang et al. [2020] provides a general deep-learning framework supporting multiple compound and protein encoders and their combinations for DTI prediction. MolTrans Huang et al. [2021] represents both drugs and proteins through learned sub-structural representations and explicitly models their interactions using a transformer-based architecture. These studies demonstrate that jointly encoding chemical and protein information is well established in DTI prediction.

More recent DTI models further employ pretrained chemical and protein representations together with attention-based interaction mechanisms. CMA-DTI Qin et al. [2026], for example, combines ChemBERTa-derived SMILES representations with ESM-2 protein embeddings and graph-based molecular representations, using attention mechanisms to model drug–protein relationships. Importantly, DTI studies have also considered *cold-drug* or unseen-drug evaluation, in which test compounds are excluded from training, providing a conceptually related setting to zero-shot ligand evaluation.

Recent work has also applied contrastive learning to large-scale virtual screening. DrugCLIP encodes protein pockets and small molecules into a shared latent space for compound retrieval Jia et al. [2026].

Despite this methodological overlap, DTI prediction and residue-level binding-site prediction address different prediction targets. DTI methods generally determine whether a protein–compound pair interacts or estimate its binding affinity, producing a pair-level prediction. In contrast, binding-site prediction requires localization of the specific protein residues associated with ligand binding. Our work therefore does not treat the combination of protein and chemical encoders or their interaction mechanism as the primary methodological novelty. Instead, we focus on systematically evaluating ligand-aware *residue-level* prediction across overrepresented, underrepresented, and previously unseen ligand regimes.

Across residue-level PLM-based approaches, common strategies for class imbalance include weighted losses Littmann et al. [2021], class-balanced focal objectives with metric-learning terms Wang et al. [2024], multi-task sharing Yuan et al. [2022], and decoder pretraining for stability in few-shot regimes Pourmirzaei et al. [2025a]. Conditioning on ligand identity through task tokens Pourmirzaei et al. [2025a] or explicit molecular representations Zhang and Xie [2023] provides ligand-specific information beyond purely *ligandagnostic* sequence models. Brief summaries of earlier traditional machine-learning, non-pretrained deep-learning, and template-based approaches are provided in Appendix A.

## Method

We systematically evaluated the impact of ligand data availability and representation on protein–ligand binding-site prediction across *overrepresented, underrepresented*, and *zero-shot* evaluation regimes. We first established an evaluation framework tailored to these categories, then used a progressive three-stage modeling suite to systematically assess the contribution of increasingly informative ligand representations. Stage 1 focuses on single-ligand prediction; Stage 2 extends the model to multiple predefined ligands through ligand-identity conditioning; and Stage 3 incorporates chemical representations to enable prediction for *zero-shot* ligands whose identities are excluded from training and validation and encountered only at test time. This progression supports a systematic comparison of modeling strategies across different ligand-frequency and data-availability regimes.

### Data Preparation and Evaluation Setup

We utilized BioLiP2 Zhang et al. [2024] as our primary dataset, a comprehensive database of biologically relevant protein–ligand binding interactions sourced from the Protein Data Bank (PDB). After applying the initial quality filters and data preprocessing steps detailed in Appendix B, we obtained 5 780 ligands and 41 327 protein– ligand samples. One underrepresented ligand (CL0) was subsequently excluded because its receptor sequences could not be partitioned into the required training, validation, and test splits after CD-HIT clustering, leaving 5 779 ligands for the original evaluation framework. To evaluate performance across different sample-availability settings, we defined three frequency-based evaluation regimes according to the number of protein–ligand samples associated with each ligand. These regimes do not imply differences in receptor diversity or receptor information content.

The following category counts describe the original dataset version after CL0 exclusion, used for Stage 1 and Stage 2. The released modified Stage 3 dataset is summarized separately in Figure 1 and Table 1.

**Table 1.** Split statistics for the released modified Stage 3 dataset used for Stage 3 evaluation and revision analyses. Ligand counts represent distinct ligand IDs; sample counts represent protein–ligand dataset rows. Original dataset split statistics are reported in Table 7 in Appendix B.

| **Category** | **Ligand IDs** | **Train samples** | **Valid samples** | **Test samples** |
|---|---|---|---|---|
| Overrepresented | 45 | 17,753 | 2,666 | 5,527 |
| Underrepresented | 121 | 1,521 | 1,571 | 1,809 |
| Zero-shot | 5,583 | 0 | 0 | 10,365 |
| **Total** | **5,749** | **19,274** | **4,237** | **17,701** |

**Figure 1.**
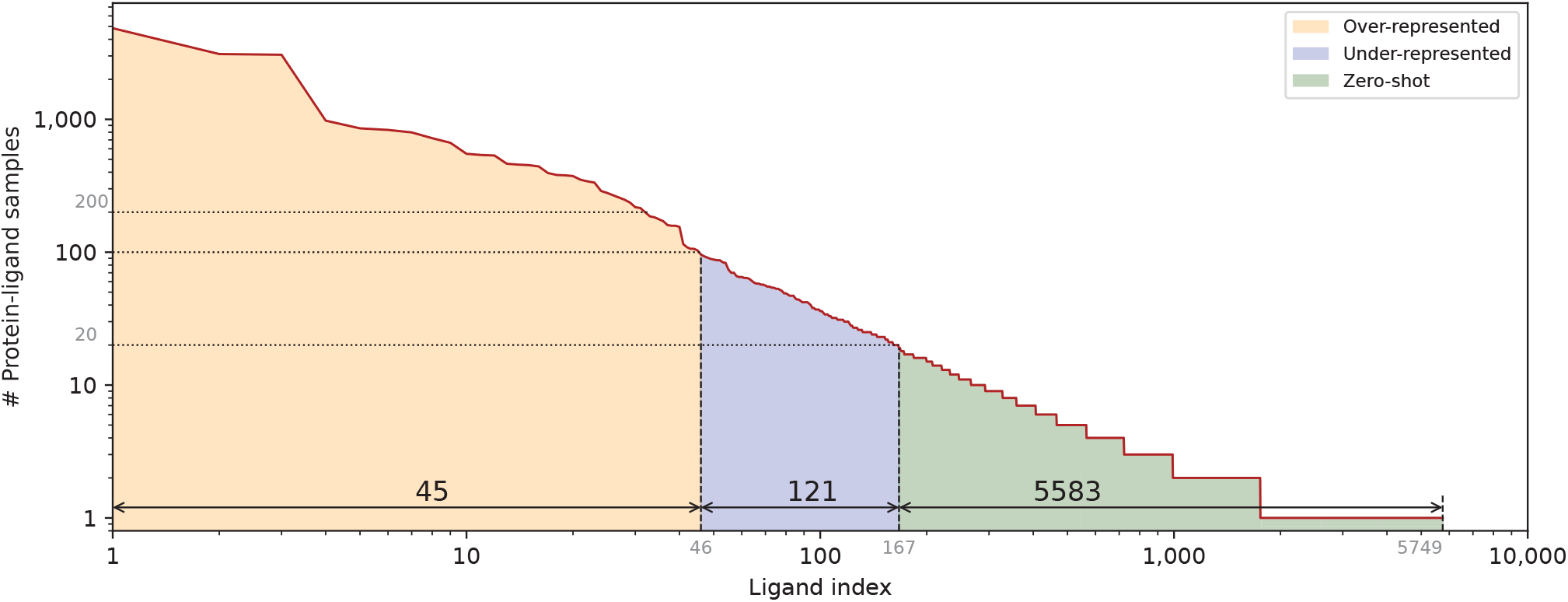
Evaluation strategies showing the distribution of ligands across three sample-count categories in the released modified Stage 3 dataset. Counts are shown on a log10 scale: 45 overrepresented, 121 underrepresented, and 5,583 zero-shot ligand IDs. Split statistics are reported in Table 1.

Overrepresented ligands (45 ligands) have at least 100 samples. Those with at least 200 samples used target training–validation–test proportions of 70%–10%–20%, whereas those with 100–199 samples used 50%–20%–30%.

Underrepresented ligands (122 ligands) have 20–99 samples. These used target training–validation–test proportions of 34%–33%–33%. *Zero-shot* ligands (5,612 ligands) have 1–19 samples and were held out entirely from training and validation, such that their ligand IDs were encountered only at test time. This regime therefore evaluates residue-level prediction for previously unseen ligand identities. Throughout this study, “zero-shot” refers to ligand identities absent from the supervised binding-site training and validation data. It does not imply unseen receptor sequences, unseen receptor families, or absence of the corresponding chemical compounds from chemical language model pretraining data.

During the original dataset construction, CD-HIT Li and Godzik [2006] clustering at a 40% sequence-identity threshold was applied within ligand-specific receptor datasets to reduce sequence redundancy when assigning protein–ligand records to training, validation, and test sets. Because clustering was performed separately within ligand-specific receptor datasets, the resulting split construction did not explicitly enforce global receptor-sequence independence across different ligands or evaluation groups. A receptor sequence, or a sequence assigned to the same global similarity cluster, may therefore occur in more than one split when associated with different ligands. Complete details of the original clustering and splitting procedure, together with an audit of exact receptor-sequence and global CD-HIT cluster overlap in the released datasets, are provided in Appendix C.

To further characterize the evaluation sets, we analyzed receptor-sequence diversity and cross-split receptor overlap using the exact modified Stage 3 datasets used for the reference evaluation. We report per-ligand receptor diversity, singleton prevalence, exact receptor-sequence overlap with training, and overlap based on global CD-HIT cluster membership. These analyses distinguish ligand-level zeroshot evaluation from receptor-level overlap and are described in Appendix C.

To further examine binding-site variability, we performed a posthoc structural analysis using available experimental reference structures associated with the modified Stage 3 test datasets. This analysis was performed only for evaluation and did not provide structural information to the prediction model. Binding-site RMSD was used to quantify structural variation among experimental reference binding sites associated with the same ligand, rather than structural error between model predictions and experimental structures. The analysis reused the existing Stage 3 predictions and involved no model retraining, new inference, threshold modification, or reconstruction of the dataset splits. Full structural mapping, quality filtering, site-matching, and RMSD procedures are provided in Appendix D.

To ensure compatibility across different CLMs, we applied minor modifications to the dataset during integration. Specifically, 29 zero-shot ligands were excluded from the final evaluation set. Details of the documented compatibility adjustments are provided in Appendix B. Stage 1 and Stage 2 used the original dataset version after CL0 exclusion. Stage 3 and the revision analyses used the previously released modified dataset summarized in Table 1, which was not changed during revision. This released version contains 5,749 ligand IDs, with 19,274 training rows, 4,237 validation rows, and 17,701 test rows. It incorporates the 29 previously reported zero-shot ligand exclusions and also excludes the underrepresented ligand F43. Its split assignments differ from the original version. Historical preprocessing logs are unavailable, so the reason for F43’s exclusion and the precise origin of the split differences could not be reconstructed.

All 5,749 evaluated ligand SMILES parsed successfully in the current RDKit audit. In the released MolFormer evaluation, chemical inputs are limited to 128 tokens, including special tokens. Three underrepresented and 72 zero-shot ligand inputs exceed this window and are truncated during evaluation. In addition, 12 evaluated zero-shot ligands contain more than 100 heavy atoms.

The modified Stage 3 evaluation groups also differed in ligand composition and sample availability. The zero-shot group contained substantially more ligand IDs, fewer samples per ligand, and a larger proportion of organic or carbon-containing ligands than the represented groups. In contrast, median heavy-atom counts and molecular weights were broadly similar across the three groups, although their observed ranges differed substantially. Detailed ligand-composition and molecular-size statistics are provided in Appendix B.1.

### Predictor Architecture

The proposed architecture leverages transformer-based models, specifically utilizing BERT-style pretrained language models tailored separately for proteins (PLMs) and chemicals (CLMs). PLMs such as ESM-2 Lin et al. [2023] provide contextualized residue-level embeddings derived solely from protein sequences. CLMs such as *MolFormer* Ross et al. [2022] encode ligand structures from SMILES strings, providing chemically meaningful representations. These pretrained encoders supply robust feature embeddings that are foundational to accurate protein–ligand binding site prediction.

#### Stage 1: Baseline Model with Single-Ligand Prediction

Stage 1 establishes our baseline architecture, which predicts binding sites for *one* ligand type at a time. Let *p* be an input protein sequence of fixed length *L*. The sequence is tokenized and passed through a *pretrained* protein language model *G*_***θ***_, whose parameters are denoted by ***θ***. This model outputs contextualized residue-level embeddings

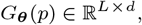

where *d* is the hidden dimension. These embeddings are fed into a linear predictor (binary classification head) *C*_***ϕ***_, parameterized by ***ϕ***, to produce a single logit for each residue (Equation 1). A separate Stage 1 model is trained for each ligand type, with its own protein encoder and classification head and using ligand-specific training, validation, and test data. Stage 1 receives only the protein sequence as input; neither the ligand SMILES representation nor a ligand identifier is provided to the model. Thus, Stage 1 performs ligand-specific prediction through separately trained models rather than through ligand conditioning within a shared model (Figure 2).

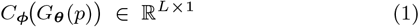

**Figure 2.**
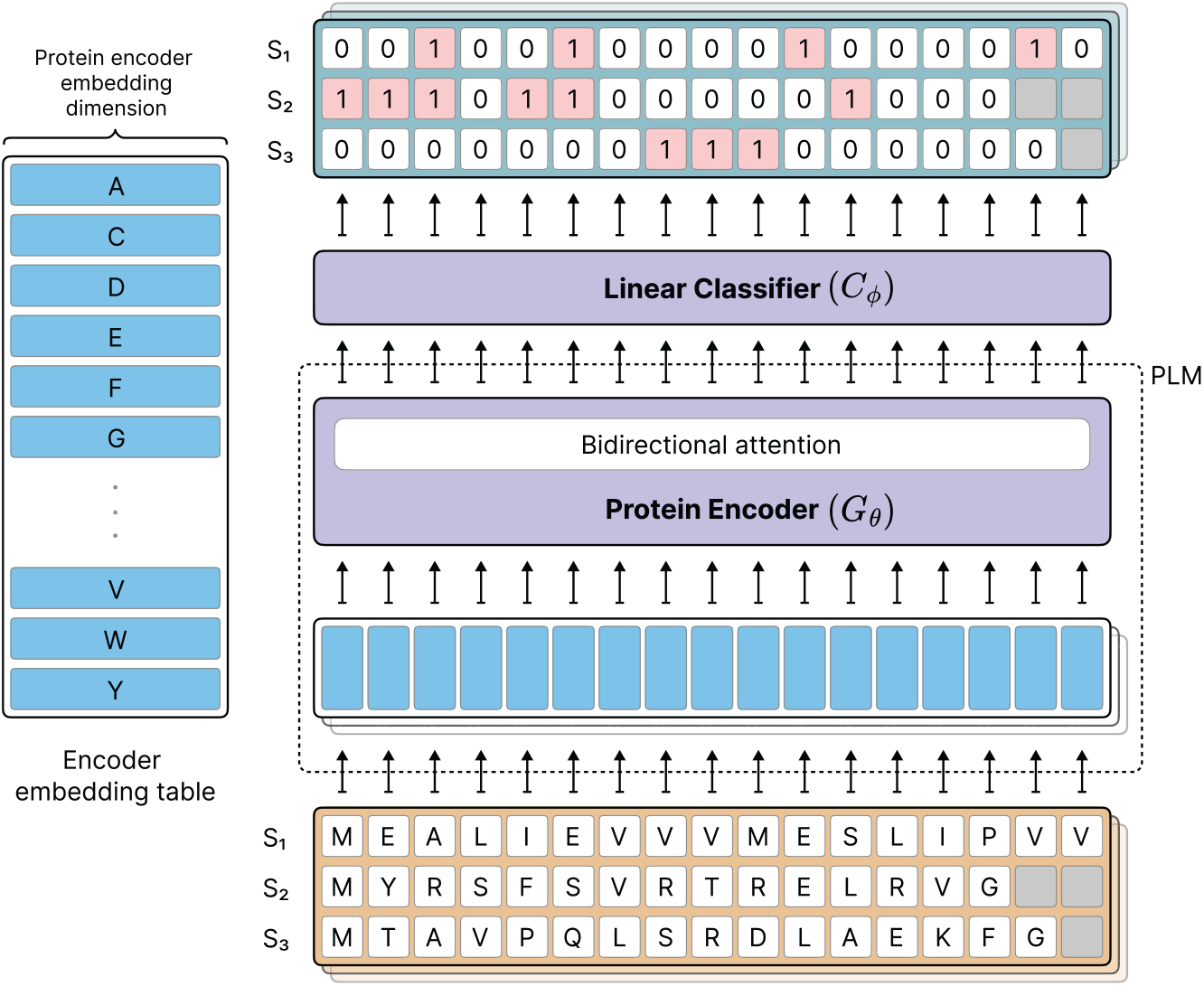
Stage 1 (single-ligand, sequence-only) baseline. A protein sequence *p* is tokenized and encoded by a pretrained PLM *G*_***θ***_, producing residue embeddings *G*_***θ***_(*p*) *∈* ℝ^*L*×*d*^. A linear head *C*_***ϕ***_ outputs residue-wise binding logits **ŷ** *∈* ℝ^*L*×1^ (Equation 1). No ligand conditioning is used; a separate model is trained per ligand type. Gray cells indicate padding or masked positions.

The Stage 1 model is trained using binary cross-entropy (BCE). In all three stages of the main experiments, padding is excluded from the training loss and evaluation metrics. The loss is averaged over valid residues within the retained sequence length. The same residue mask is used for checkpoint selection and final evaluation.

For batch *t*, the loss is computed as

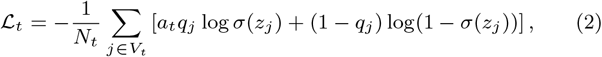

where *V*_*t*_ is the set of valid residue positions, excluding padding, and *N*_*t*_ = |*V*_*t*_| is the number of these positions. The quantity *z*_*j*_ is the output logit for position *j*, and *σ*(*z*_*j*_) is the corresponding sigmoid probability. When label smoothing is disabled, *q*_*j*_ = *y*_*j*_, where *y*_*j*_ *∈* {0, 1} is the binary binding label. The batch-dependent coefficient *a*_*t*_ weights only the positive-label term; no separate dataset-level multiplier or residue-specific learned weight is used.

To account for class imbalance, let

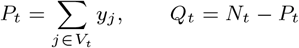

denote the numbers of positive and negative valid residue positions in batch *t*, respectively.

The batch ratio is defined as

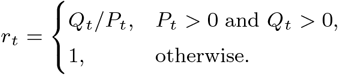

At the beginning of each training epoch or validation/evaluation pass, the weight state is reset, and the first batch uses *a*_1_ = *r*_1_. For subsequent batches, the coefficient is updated according to

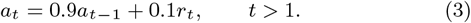

The positive-class weight is updated using an exponential moving average of the negative-to-positive residue ratio with smoothing coefficient 0.9. Counts exclude padding. The weight is initialized from the first batch and is not capped at 1. The moving average reduces batch-to-batch fluctuations while retaining positive-class upweighting.

#### Stage 2: Multi-Ligand Prediction with Ligand-Specific Conditioning

Stage 2 augments the baseline by conditioning on ligand identity, enabling simultaneous prediction of binding sites for multiple *predefined* ligand types. Protein sequences *p* are encoded by the same PLM *G*_***θ***_ used in Stage 1, producing residue embeddings

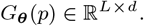

Ligand information is provided as an index *ℓ* that queries a trainable embedding table *E*_***η***_, parameterized by ***η***, yielding a ligand embedding

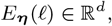

analogous to word embeddings (e.g., word2vec Church [2017]). A bidirectional transformer decoder *T*_***ψ***_, with parameters ***ψ***, integrates protein and ligand representations via cross-attention. Similar to Stage 1, a linear classification head *C*_***ϕ***_ produces residue-level logits (Equation 4). This stage supports all ligand types seen during training, but cannot generalize to unseen ligands (Figure 3).

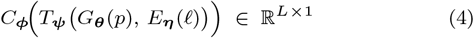

**Figure 3.**
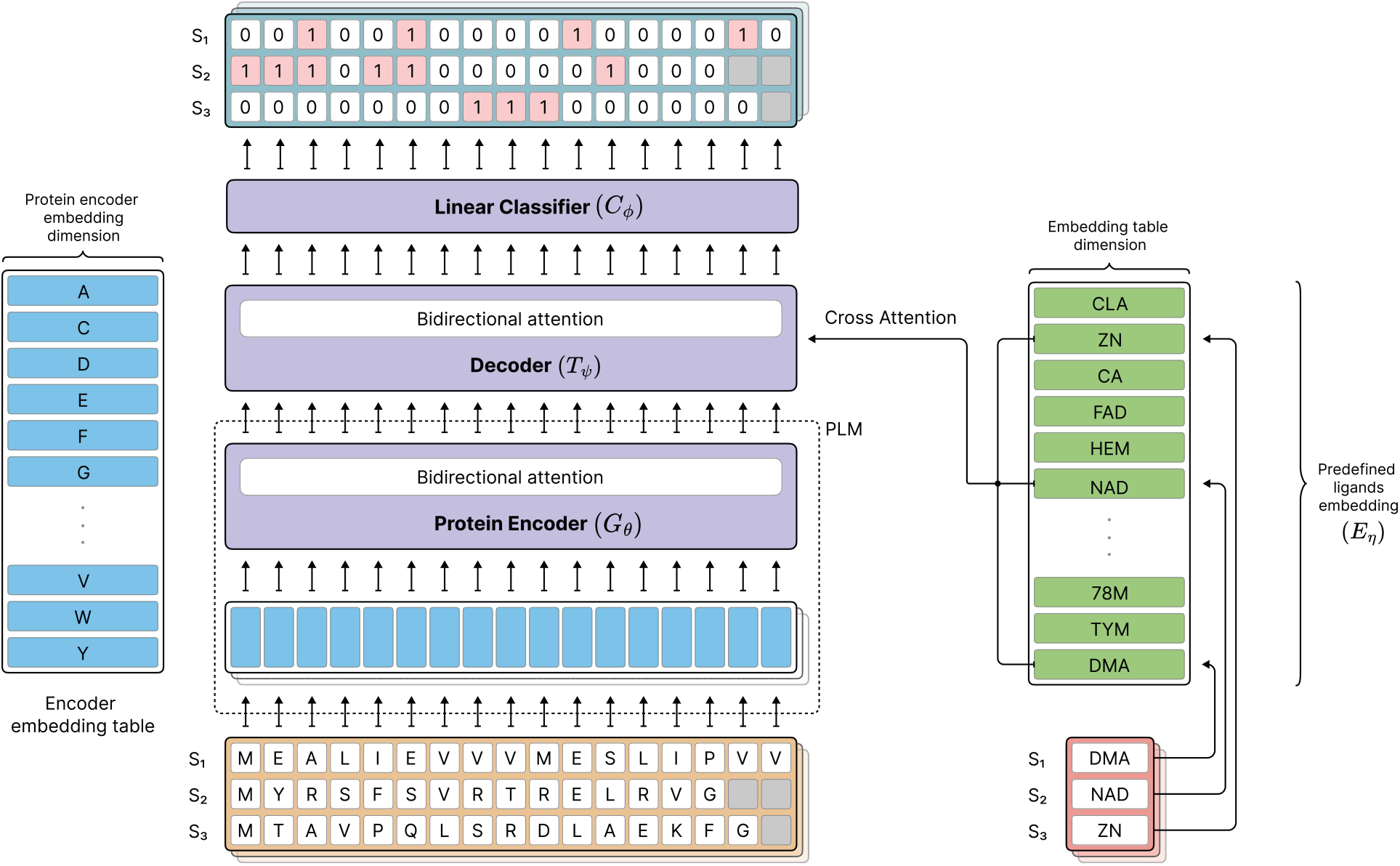
Stage 2 (multi-ligand with learned ligand embedding). Protein embeddings *G*_***θ***_(*p*) *∈* ℝ^*L*×*d*^ are produced as in Stage 1. Ligand identity *ℓ* indexes a trainable embedding table *E*_***η***_(*ℓ*) *∈* ℝ^*d*^. A bidirectional decoder *T*_***ψ***_ fuses protein and ligand features via cross-attention, and the classifier *C*_***ϕ***_ outputs residue-wise logits (Equation 4). The model supports all ligands observed during training but does not generalize to novel ligands. Gray cells indicate padding or masked positions.

#### Stage 3: Zero-Shot Generalization with a Pretrained Chemical Language Model

Stage 3 equips the model with *zero-shot* capability by replacing the fixed ligand-embedding mechanism with a pretrained chemical language model. Protein sequences *p* are encoded by the same PLM *G*_***θ***_ as in previous stages. Each ligand is supplied directly as a SMILES string *s* and embedded by the chemical language model *F*_***ω***_, whose parameters are denoted by ***ω***, yielding an embedding

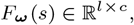

where *l* is the SMILES token length and *c* the CLM feature dimension, typically distinct from the protein feature dimension *d*.

To align the feature dimensions of the two modalities, we introduce a learnable linear projector

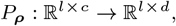

parameterized by ***ρ***, producing projected ligand embeddings

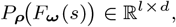

which preserve the SMILES token length *l* while mapping the CLM feature dimension from *c* to the protein feature dimension *d* for cross-attention within the decoder *T*_***ψ***_. The fused representations are passed to the binary classifier head *C*_***ϕ***_. By using SMILES-derived representations rather than a fixed ligand-identity embedding table, the architecture (Equation 5) supports residue-level prediction for ligand identities absent from the supervised binding-site training and validation data (Figure 4). This capability does not establish that these compounds were absent from the CLM pretraining corpus.

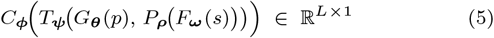

**Figure 4.**
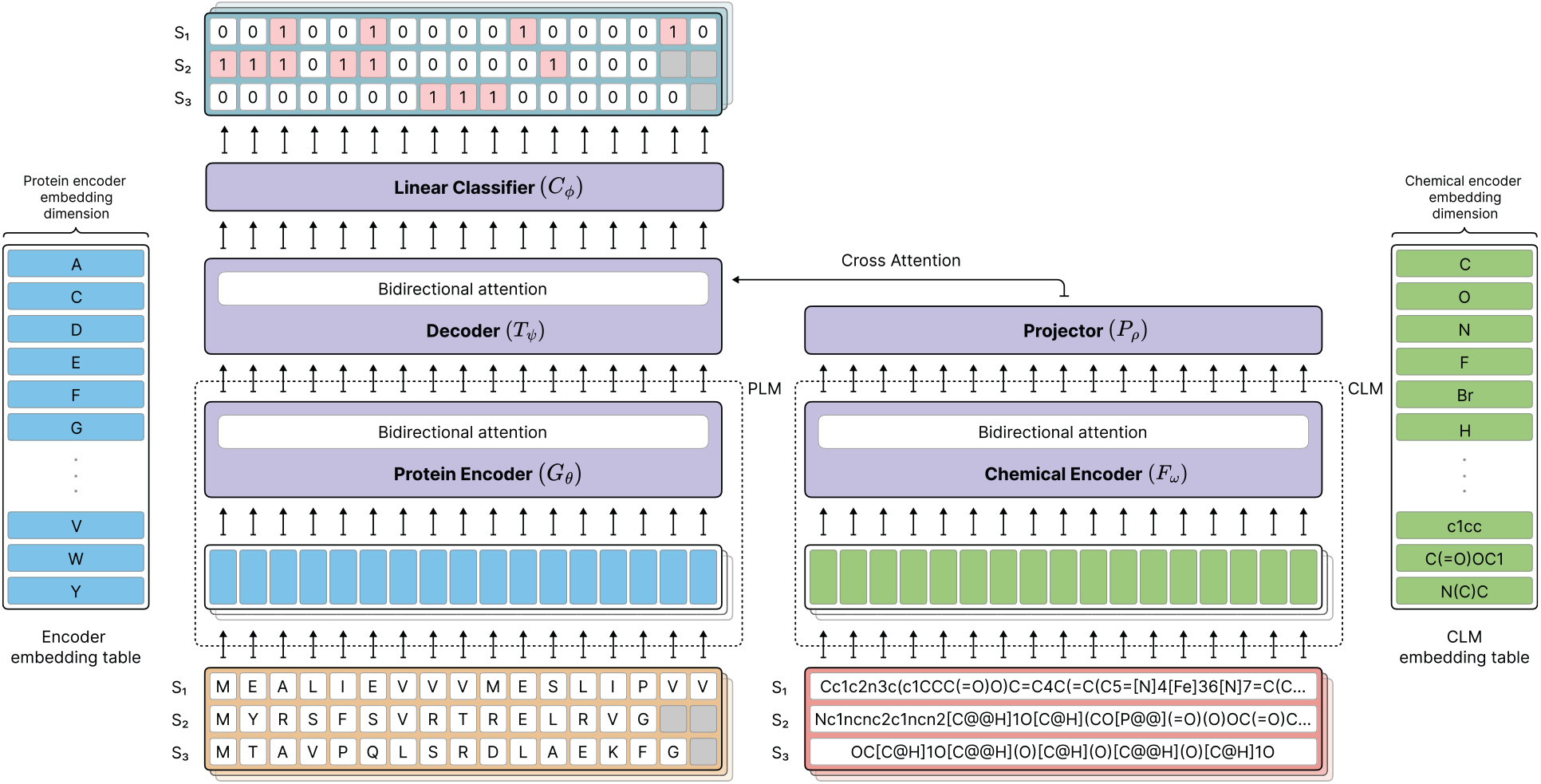
Stage-3 (*zero-shot*) architecture. A protein sequence *p* of length *L* is embedded by a pretrained PLM *G*_***θ***_ (left), and the ligand SMILES *s* with *l* tokens is embedded by a pretrained CLM *F*_***ω***_ (right). A learnable projector *P*_***ρ***_ maps CLM features to the protein feature dimension, and a bidirectional decoder *T*_***ψ***_ fuses the two streams via cross-attention; a linear head *C*_***ϕ***_ then outputs residue-wise binding logits **ŷ** *∈* ℝ^*L*×1^ (Equation 5). Gray cells indicate padding/masks. Conditioning on SMILES rather than a fixed ligand ID enables inference on unseen ligands (*zero-shot* residue-level prediction).

### Evaluation Metrics

We evaluate residue-level binary predictions primarily using precision, recall, pooled *F*_1_, ligand-macro *F*_1_, and Matthews correlation coefficient (MCC). Because binding residues constitute a small fraction of all protein residues, overall accuracy can be dominated by the majority non-binding class and is therefore not used as a primary measure of predictive performance.

Let *y*_*i*_ *∈* {0, 1} denote the ground-truth binding label for residue *i* and *ŷ*_*i*_ *∈* [0, 1] the predicted binding probability. A hard prediction is obtained with a threshold *τ* (default *τ* = 0.5):

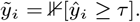

Over a set of valid residues, excluding padding, we compute the counts TP, FP, TN, and FN.

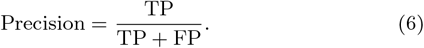

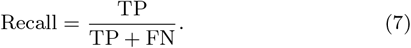

The *F*_1_ score is the harmonic mean of precision and recall:

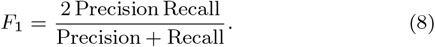

When reported as pooled *F*_1_, the counts of true positives, false positives, and false negatives are combined across all valid residues before computing the score.

For a ligand group L, we compute 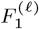 separately for each ligand *ℓ* and report the unweighted average:

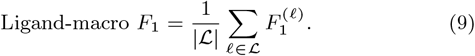

Thus, pooled *F*_1_ weights ligands according to their contributions to the aggregate residue counts, whereas ligand-macro *F*_1_ gives each ligand equal weight.

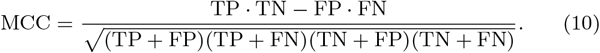

Unless otherwise noted, all metrics are computed at the residue level on held-out test sets using the default threshold *τ* = 0.5.

## Experiments

We constructed a base architecture and systematically extended it in two progressive stages to enable comprehensive evaluation across all ligand representation categories. These categories include *over-represented, underrepresented*, and *zero-shot* ligands. Across all experiments, we evaluated multiple combinations of pretrained language models: protein language models (PLMs) including several scales of ESM-2, and chemical language models (CLMs) including *MolFormer* and *UniMol-2* Ji et al. [2024].

Optimization employed the AdamW optimizer Loshchilov [2017] with a weight decay of 0.01, momentum parameters *β*_1_ = 0.9 and *β*_2_ = 0.98, and *ϵ* = 10^−7^. We used a cosine-annealing learning-rate schedule Loshchilov and Hutter [2016] without warm-up, starting at 5 × 10^−5^ and decaying to zero over the training horizon. All models were implemented in PyTorch 2.6.0 Ansel et al. [2024] and trained using mixed-precision BF16 arithmetic Kalamkar et al. [2019], Micikevicius et al. [2017] on an NVIDIA A100 GPU with 80 GB of memory. Detailed hyperparameter settings and additional implementation notes are provided in Appendix H. The reference Stage 3 configuration is summarized in Appendix H (Table 18).

### Stage 1: Single-Ligand Prediction

In Stage 1, we trained individual models for each ligand type without using any ligand-specific embeddings or chemical representations. This stage was restricted to the 45 *overrepresented* ligands in our dataset, and each ligand was modeled independently using identical hyperparameters. All experiments in this stage used the ESM-2 650M model as the protein encoder.

For every ligand, training was monitored using the corresponding validation split, and the best-performing checkpoint was selected based on the peak validation *F*_1_ score. Final results were then computed on the held-out *overrepresented* test set. Each ligand’s results were obtained from a single training run. To characterize seed-to-seed variability, we additionally retrained a representative six-ligand subset using three random seeds as part of the matched linear-predictor–decoder ablation. Across these Stage 1 linear-predictor runs, the per-ligand standard deviations in *F*_1_ ranged from 0.0005 to 0.0281. Full per-ligand results and experimental details are provided in Appendix F.

The overall performance across ligands is summarized in Tables 2 and 4, with additional aggregated statistics provided in Table 20. These results establish a detailed residue-level binary classification baseline for subsequent stages.

**Table 2.** Benchmarking the three stages on the test sets. Ligand-macro *F*_1_ is computed by calculating *F*_1_ separately for each ligand type and averaging across ligand types, thereby giving each ligand equal weight. The reported *F*_1_ is the pooled residue-level *F*_1_ score. For Stage 2 and Stage 3, values are reported as mean ± standard deviation across three independent training seeds.

| Method | Training Dataset | Overrepresented | Underrepresented | Zero-shot |  |
|---|---|---|---|---|---|
|  |  | Ligand-macro *F*<sub>1</sub> | Ligand-macro *F*<sub>1</sub> | Ligand-macro *F*<sub>1</sub> | **Pooled** *F*<sub>1</sub> |
| Stage 1 | Overrepresented (separated) | 0.4769 | – | – | – |
| Stage 2 | Overrepresented | 0.5826 ± 0.0035 | – | – | – |
| Stage 2 | Overrepresented + Underrepresented | 0.5832 ± 0.0014 | 0.3752 ± 0.0049 | – | – |
| Stage 3 | Overrepresented + Underrepresented | 0.5526 ± 0.0012 | 0.3603 ± 0.0029 | 0.2338 ± 0.0051 | 0.3109 ± 0.0087 |

**Table 3.**
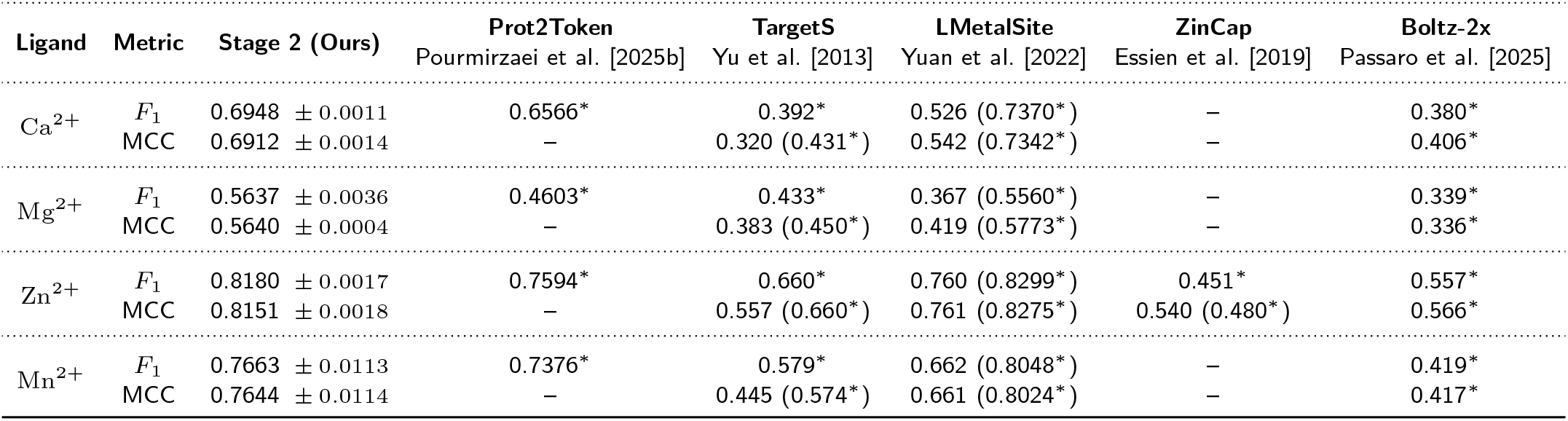
Comparison of Stage 2 performance with other available methods on four selected metal ions in the *overrepresented* test set based on *F*_1_ scores. These four metal ions were selected for their biological importance and the availability of published binding-site prediction results, enabling comparison with multiple baseline methods. Main baseline values are taken from the original papers; ^*\**^ indicates methods re-evaluated on our test sets.

| **Ligand** | **Metric** | **Stage 2 (Ours)** | **Prot2Token**<br>Pourmirzaei et al. [2025b] | **TargetS**<br>Yu et al. [2013] | **LMetalSite**<br>Yuan et al. [2022] | **ZinCap**<br>Essien et al. [2019] | **Boltz-2x**<br>Passaro et al. [2025] |
|---|---|---|---|---|---|---|---|
| Ca<sup>2+</sup> | *F*<sub>1</sub> | 0.6948 ± 0.0011 | 0.6566<sup>*</sup> | 0.392<sup>*</sup> | 0.526 (0.7370<sup>*</sup>) | – | 0.380<sup>*</sup> |
|  | MCC | 0.6912 ± 0.0014 | – | 0.320 (0.431<sup>*</sup>) | 0.542 (0.7342<sup>*</sup>) | – | 0.406<sup>*</sup> |
| Mg<sup>2+</sup> | *F*<sub>1</sub> | 0.5637 ± 0.0036 | 0.4603<sup>*</sup> | 0.433<sup>*</sup> | 0.367 (0.5560<sup>*</sup>) | – | 0.339<sup>*</sup> |
|  | MCC | 0.5640 ± 0.0004 | – | 0.383 (0.450<sup>*</sup>) | 0.419 (0.5773<sup>*</sup>) | – | 0.336<sup>*</sup> |
| Zn<sup>2+</sup> | *F*<sub>1</sub> | 0.8180 ± 0.0017 | 0.7594<sup>*</sup> | 0.660<sup>*</sup> | 0.760 (0.8299<sup>*</sup>) | 0.451<sup>*</sup> | 0.557<sup>*</sup> |
|  | MCC | 0.8151 ± 0.0018 | – | 0.557 (0.660<sup>*</sup>) | 0.761 (0.8275<sup>*</sup>) | 0.540 (0.480<sup>*</sup>) | 0.566<sup>*</sup> |
| Mn<sup>2+</sup> | *F*<sub>1</sub> | 0.7663 ± 0.0113 | 0.7376<sup>*</sup> | 0.579<sup>*</sup> | 0.662 (0.8048<sup>*</sup>) | – | 0.419<sup>*</sup> |
|  | MCC | 0.7644 ± 0.0114 | – | 0.445 (0.574<sup>*</sup>) | 0.661 (0.8024<sup>*</sup>) | – | 0.417<sup>*</sup> |

**Table 4.**
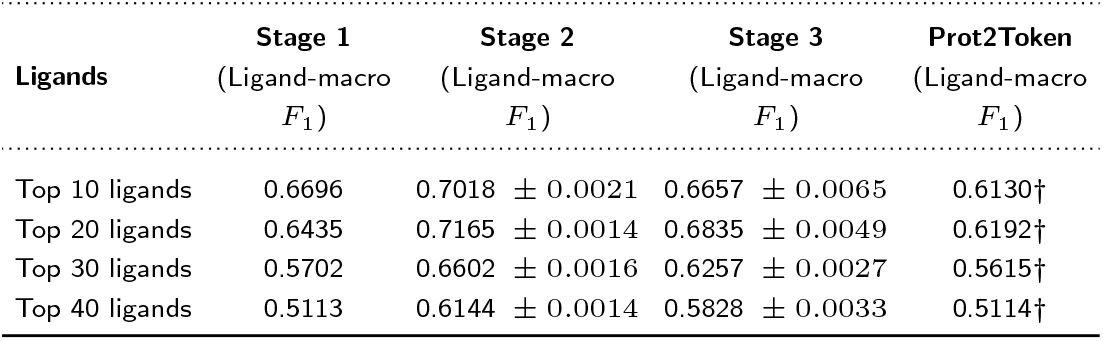
Comparison of Prot2Token with our approach on the top 40 ligands in the *overrepresented* test set. †: values are reported from the original paper.

Despite the uniform training procedure, we observed considerable variation in model performance across ligand types. Several ligands, such as `FAD` and `ZN`, achieved high *F*_1_ scores, whereas other ligands—including `BMA` and `MAN`—exhibited near-zero predictive performance. While some of these discrepancies may reflect differences in sample size and receptor composition, the observed variation motivates further analysis of receptor and binding-site variability across ligands.

### Stage 2: Multiple Ligands

In Stage 2, we evaluated the impact of extending the architecture to support multiple ligand types via ligand-specific conditioning. The introduction of a trainable ligand-embedding table enabled a single model to learn from a pooled dataset spanning many ligand classes, in contrast to the per-ligand training of Stage 1. We assessed this architecture under two training regimes: (i) training on only the 45 *overrepresented* ligands used in Stage 1, and (ii) training on an expanded dataset that combined both *overrepresented* and *underrepresented* ligands. For both regimes, reported results are presented as mean ± standard deviation across three independent training seeds (Table 2).

In both settings, the Stage 2 model achieved substantially stronger predictive performance than the single-ligand Stage 1 baseline (Table 20). However, the original Stage 1–Stage 2 comparison simultaneously changes predictor architecture, ligand conditioning, and the training regime, and therefore does not isolate the source of this improvement. In a matched six-ligand ablation that retained ligand-specific training, replacing the Stage 1 linear predictor with a Stage 2-compatible decoder produced only a modest average improvement in ligand-macro *F*_1_ (0.5431±0.0048 to 0.5567±0.0198; paired difference +0.0136 ± 0.0171). This suggests that decoder architecture alone is unlikely to account for the larger Stage 1–Stage 2 performance difference. Full per-ligand results, three-seed variability, and matched training settings are provided in Appendix F. A comparison with Prot2Token on the *overrepresented* test set appears in Table 4.

As in Stage 1, all experiments in this stage used the ESM-2 650M protein language model (hyperparameters summarized in Table 17). Fair benchmarking across sequence- and structure-based predictors is complicated by potential data contamination, since many structure-based systems are pretrained or tuned on PDB-derived sequences or complexes that may partially overlap with evaluation data through homology. Although our dataset construction reduces within-ligand sequence redundancy through CD-HIT clustering at 40% sequence identity (Appendix B), cross-corpus contamination remains a general concern when comparing to external tools.

To provide a strong structure-based baseline under controlled conditions, we include Boltz-2x, the Boltz-2 configuration with physicality steering potentials enabled Passaro et al. [2025], a recent high-performing open-source protein–ligand complex predictor. We ran Boltz-2x without multiple sequence alignments (MSAs) and truncated protein sequences to length 1280, supplying only single protein sequences as input. These restrictions remove the advantage of explicit MSA input, which is not supplied to our model. To convert Boltz-2x predictions into residue-level binding-site labels, we enumerated intermolecular atomic contacts: a contact is defined as any protein–ligand atom pair within a distance less than or equal to the sum of their van der Waals radii plus 0.5 Å. A residue is labeled as binding if it forms more than two such contacts. This yields residue-level labels directly comparable to our model outputs.

### Stage 3: Zero-Shot Ligands

In Stage 3, we assessed the impact of replacing the ligand-embedding table from Stage 2 with a pretrained chemical language model (MolFormer), enabling ligand conditioning directly from SMILES strings. This representation supports evaluation on ligand identities absent from the supervised binding-site training and validation data, without requiring a learned identity embedding for each test ligand.

The model was trained using the combined *overrepresented* and *underrepresented* training sets. Evaluation was performed on all three test groups—*overrepresented, underrepresented*, and *zero-shot* ligands—with results summarized in Table 2. The principal Stage 3 results in Table 2 are reported as mean ± standard deviation across three independent training seeds to assess seed-to-seed variability.

We additionally include a direct comparison to Prot2Token Pourmirzaei et al. [2025b] on the *overrepresented* test set (Table 4). Prot2Token serves as an important reference point because it is among the strongest published sequence-only binding-site predictors, outperforming multiple competing deep learning methods. For fairness, our train/validation/test splits are constructed to closely match those used in Prot2Token, using the same BioLiP2 source data, with differences arising only from minor ligand-specific sample-count variations. This alignment ensures that comparisons across methods reflect architectural differences rather than dataset inconsistencies.

When evaluating with the same set of hyperparameters used for the prior stages, we found that the Stage 3 model underperformed on both the *overrepresented* (0.5526 vs. 0.5832 ligand-macro *F*_1_) and *underrepresented* (0.3603 vs. 0.3752 ligand-macro *F*_1_) test sets compared to Stage 2.

Nevertheless, Stage 3 offered a distinct advantage in its ability to perform *zero-shot* inference, achieving a pooled residue-level *F*_1_ of 0.3109 on the test set of previously unseen ligands.

Singleton ligands constituted 71.7% of zero-shot ligand IDs. Performance for singleton and non-singleton ligands is reported separately in Appendix C (Table 11) to aid interpretation of ligand-macro *F*_1_.

Zero-shot here refers to ligand IDs absent from supervised training and validation, rather than a guarantee of chemical novelty. A chemical-input audit found 31 zero-shot IDs with matching stereoaware canonical input representations in training. Excluding these matches had a very small effect on single-checkpoint pooled residue-level *F*_1_, changing it from 0.3193 to 0.3179. Per-ligand *F*_1_ had a weak positive association with similarity to the nearest training ligand. The chemical-overlap analysis and similarity-stratified results are reported in Appendix C.4.

To further characterize Stage 3 performance under class imbalance, we report precision and recall for the released Stage 3 checkpoint in Table 5. These results are from a single checkpoint and are therefore reported separately from the three-run mean ± standard deviation results in Table 2.

**Table 5.**
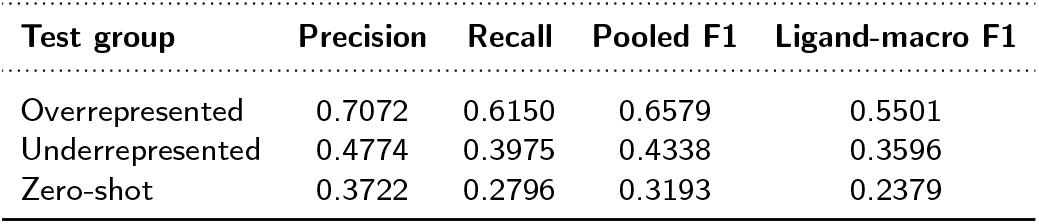
Precision, recall, pooled F1, and ligand-macro F1 for the released Stage 3 checkpoint using native ligand inputs. These single-checkpoint results are reported separately from the three-run mean ± standard deviation results in Table 2.

For the zero-shot group, precision (0.3722) exceeded recall (0.2796), indicating that missed binding residues contributed substantially to the observed F1 performance.

### Ligand-input control experiments

To determine whether Stage 3 uses ligand-specific information from the SMILES input rather than relying solely on protein-derived features or generic ligandability, we performed three control experiments using a single trained Stage 3 checkpoint. These controls were applied only at inference time; the model was not retrained. The native ligand SMILES were replaced with (i) a constant SMILES input shared within each test group, (ii) mismatched SMILES obtained by randomly reassigning ligand inputs across protein–ligand pairs, or (iii) scrambled SMILES generated by randomly permuting the characters of each ligand SMILES as an input-corruption control.

For each native-versus-control comparison, uncertainty in the pooled *F*_1_ difference was estimated using 2,000 paired bootstrap resamples of exact receptor-sequence groups within each test set. Native and control observations were resampled together.

As shown in Table 6, using the native ligand SMILES produced the highest pooled *F*_1_ score in all three evaluation groups. For the overrepresented group, the native-SMILES *F*_1_ score was 0.6579, compared with 0.3610, 0.2622, and 0.5357 for the constant, mismatched, and scrambled controls, respectively. For the underrepresented group, the corresponding scores were 0.4338, 0.3381, 0.2799, and 0.3210. In the zero-shot group, the native-SMILES score was 0.3193, compared with 0.1335, 0.2393, and 0.2583 for the three controls. Across all nine native-versus-control comparisons, the 95% bootstrap confidence intervals for the native-minus-control pooled *F*_1_ differences were above zero (Table 16). These results show that preserving the correct ligand input yields higher pooled *F*_1_ than each control across all three evaluation regimes, supporting a contribution from the supplied ligand association to Stage 3 predictions.

**Table 6.** Ligand-input control experiments for the trained Stage 3 model. Control inputs were introduced only at inference time without retraining. Native denotes each protein sequence paired with the correct bound-ligand SMILES. Values report pooled F1 scores. The native-SMILES values correspond to this single checkpoint and therefore may differ from the three-run mean results reported in Table 2.

| **Test group** | **Native** | **Constant** | **Mismatched** | **Scrambled** |
|---|---|---|---|---|
| Overrepresented | **0.6579** | 0.3610 | 0.2622 | 0.5357 |
| Underrepresented | **0.4338** | 0.3381 | 0.2799 | 0.3210 |
| Zero-shot | **0.3193** | 0.1335 | 0.2393 | 0.2583 |

### Binding-Site Structural Variability Analysis

To examine whether prediction performance was associated with structural variability among experimental reference binding sites, we compared per-ligand covered F1 with binding-site RMSD in the structurally covered subset. The raw Spearman correlations were 0.299, 0.244, and 0.168 for the overrepresented, underrepresented, and zero-shot groups, respectively, and decreased to 0.043, 0.023, and 0.038 after adjustment. Thus, within the structurally covered subset, better performance was not consistently associated with greater conservation of the experimental reference binding sites. Full methods, coverage information, and structural analyses are provided in Appendix D.

Importantly, structural coverage selected a higher-performing subset of the reference population. For the released Stage 3 checkpoint, pooled F1 increased from 0.6579 to 0.7463 for the overrepresented group, from 0.4338 to 0.5677 for the underrepresented group, and from 0.3193 to 0.4044 for the zero-shot group when evaluation was restricted to the structurally covered samples. Valid structural comparisons covered only 18.6%, 30.0%, and 18.6% of samples in the overrepresented, underrepresented, and zero-shot groups, respectively. Thus, the structural analyses characterize a selected, higher-performing subset of the test population, and their findings should not be generalized directly to the complete test sets. In particular, these analyses do not rule out memorization outside the structurally covered subset (Fig. 12).

We additionally examined binding-site sequence conservation within the same structurally covered subset by relating reference-site amino-acid identity to per-ligand covered F1. The Spearman correlations between site sequence identity and covered F1 were −0.258, −0.086, and −0.128 for the overrepresented, underrepresented, and zero-shot groups, respectively. After adjustment, the corresponding correlations were −0.087, 0.115, and 0.095. Thus, binding-site amino-acid identity did not show a stable positive relationship with predictive performance, and the direction of the association changed after adjustment in two groups. These results do not establish that conservation is harmful or rule out memorization of conserved binding sites (Fig. 13).

### Scaling comparison

To better understand the impact of model scale on protein–ligand binding site prediction, we conducted a series of ablation experiments varying both the size of the PLM backbone and the choice of CLM on the Stage 3 architecture. Specifically, we evaluated model performance across five ESM-2 sizes (8M, 35M, 150M, 650M, and 3B), paired with different CLM configurations to assess how scaling affects predictive capability. In addition to varying the backbone size, we compared multiple CLMs, including MolFormer, UniMol-2 (both 84M and 570M parameter variants), and partially unfrozen versions of the 84M UniMol-2 model.

Across all three stages, the final three transformer blocks of ESM-2 were fine-tuned, while the remaining ESM-2 blocks and embedding layer were kept frozen. For the Stage 3 scaling experiments, MolFormer was kept fully frozen. UniMol-2 was also kept fully frozen in the standard 84M and 570M configurations. In two additional UniMol-2 84M variants, the final two or four transformer blocks were unfrozen and fine-tuned, respectively, to assess the effect of partial CLM fine-tuning.

To ensure a fair comparison across architectures, we held the training recipe fixed: similar optimization hyperparameters and the same architectural configurations (Tables 17 and 19). Each configuration was trained with three random seeds, and we report mean results. However, the individual per-seed results for Figure 5 are no longer available, preventing reconstruction of standard deviations for these scaling comparisons. As shown in Figure 5, increasing the protein encoder size generally improves pooled residue-level *F*_1_ across the evaluated test sets, although the trend is not strictly monotonic across all configurations. In contrast, increasing the chemical encoder scale (e.g., UniMol-2 84M→570M) produces smaller and less consistent gains. These observations apply to the tested model configurations and training settings and should not be interpreted as a general scaling relationship for protein and chemical encoders.

**Figure 5.**
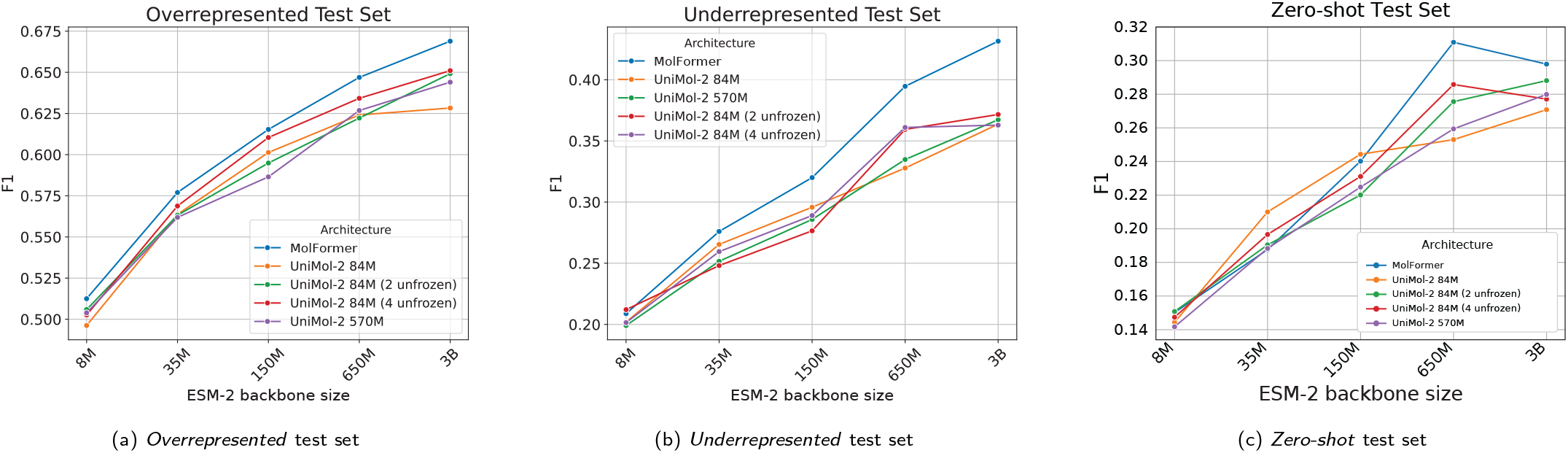
Results of ablation studies on the three ligand test sets based on pooled residue-level *F*_1_: (a) *overrepresented*, (b) *underrepresented*, and (c) *zero-shot* test sets. The numerical values underlying the curves are reported in Table 22.

## Discussion

In this study, we systematically evaluated residue-level protein– ligand binding site prediction across three ligand-frequency regimes: *overrepresented, underrepresented*, and *zero-shot* ligands. Using the progressive three-stage modeling suite, we examined how prediction performance changes as the available ligand information progresses from ligand-specific training (Stage 1), to ligand-identity conditioning within a shared model (Stage 2), and finally to chemical representations derived from SMILES (Stage 3). This evaluation reveals distinct performance patterns across the three regimes. In the original comparison, Stage 2 achieved a higher ligand-macro *F*_1_ than Stage 1 on the overrepresented test set (0.5832 versus 0.4769; Table 2). However, this comparison simultaneously changes predictor architecture, ligand conditioning, and the training regime. In the matched six-ligand ablation retaining ligand-specific training, replacing the linear predictor with a Stage 2-compatible decoder yielded a modest average change in ligand-macro *F*_1_ (paired difference +0.0136 ± 0.0171 across three seeds). This limited comparison does not isolate the contributions of ligand conditioning or pooled multi-ligand training to the overall Stage 1–Stage 2 difference. Stage 2 was also competitive with or better than the evaluated baselines on several ions and cofactors (e.g., Zn^2+^ *F*_1_ = 0.818 vs. 0.557 for Boltz-2x under residue-label conversion; Table 3). Our method predicts candidate binding residues from protein sequences and ligand SMILES without generating a 3D complex. These predictions could help guide subsequent docking or structure prediction. The current evaluation measures binding-site localization and does not establish binder discrimination or affinity-ranking performance.

Replacing the lookup embedding with a CLM over SMILES (Stage 3) extends the evaluation to *zero-shot* ligands whose identities are excluded from training and validation: on this held-out ligand set, the model reaches a pooled residue-level *F*_1_ of 0.3109, while the unweighted mean of per-ligand *F*_1_ scores (ligand-macro *F*_1_) is 0.2338 (Table 2). This zero-shot capability is accompanied by a modest decrease in performance on represented ligands relative to Stage 2 (*overrepresented* ligand-macro *F*_1_: 0.5832 to 0.5526; *underrepresented* ligand-macro *F*_1_: 0.3752 to 0.3603). Together with the ablation in Fig. 5, these results show that increasing protein-encoder (ESM-2) capacity generally improves performance across the evaluated regimes, although the trend is not strictly monotonic. In particular, for the zero-shot MolFormer setting, pooled residue-level *F*_1_ reaches 0.3109 with ESM-2 650M and decreases to 0.2979 with ESM-2 3B. In contrast, scaling the CLM produces comparatively small or inconsistent changes, suggesting that protein-representation capacity has a stronger effect on performance than chemical-encoder scale under the current architecture.

The ligand-input control experiments further clarify the contribution of ligand information in Stage 3. Replacing the native ligand SMILES with constant, mismatched, or character-scrambled inputs reduced predictive performance across all three evaluation regimes. These results support the conclusion that Stage 3 predictions depend in part on the supplied ligand association rather than solely on protein-derived or generic ligandability signals. However, these controls do not establish that the model has learned a complete chemical representation of ligand–protein interactions. In particular, the scrambled-SMILES control may also reflect sensitivity to corrupted SMILES syntax. This finding complements the CLM-scaling ablation: the limited benefit from increasing CLM scale should not be interpreted as evidence that ligand input is uninformative; rather, the control experiments show that preserving the native ligand input is beneficial under the current architecture.

A single-checkpoint subgroup analysis found that zero-shot pooled residue-level *F*_1_ did not decrease for receptors without an exact training-sequence match or without a shared 40% CD-HIT audit cluster (Appendix C.3). These descriptive results suggest that useful predictions are not restricted to training-overlapping receptors, although they do not establish an absence of memorization.

Two factors likely limit headroom. First, the strong class imbalance—reflected by very high overall accuracy despite substantially lower *F*_1_—makes optimization sensitive to calibration and decision thresholds, especially for sugars and small polar ligands where Stage 1 underperforms (e.g., `BMA, MAN`; Table 20). Second, although the ligand-input controls indicate that Stage 3 predictions depend in part on the supplied ligand association, CLM–PLM alignment is learned through a shallow projector. Stronger cross-modal fusion may therefore enable more effective integration of ligand representations with protein context, potentially recovering the small Stage 3 performance degradation on overrepresented and underrepresented ligands while further improving *zero-shot* generalization.

## Conclusion

We established a systematic ligand-aware benchmark and evaluation framework for residue-level protein–ligand binding-site prediction across *overrepresented, underrepresented*, and *zero-shot* ligand regimes. Using a progressive three-stage modeling suite, we characterized how predictive performance changes as ligand supervision decreases and demonstrated residue-level prediction for previously unseen ligands using protein sequences and SMILES alone. Importantly, ligand-input control experiments using constant, mismatched, and scrambled SMILES reduced predictive performance across all three evaluation regimes, supporting a contribution from the supplied ligand information to Stage 3 predictions rather than reliance solely on protein-derived or generic ligandability signals. Together, these results provide a systematic benchmark and evaluation framework for studying ligand-aware residue-level prediction under varying levels of ligand data availability.

Predicted binding residues could help guide subsequent docking or structure prediction. Binder discrimination and affinity ranking remain to be evaluated. We release code, data, and model weights to support reproducible progress in ligand-aware and zero-shot residue-level binding prediction: https://github.com/mahdip72/ProteinLigand.

## Code & Data Availability

The source code used in this study is publicly available at https://github.com/mahdip72/ProteinLigand. The original datasets and the modified datasets used for the revised Stage 3 evaluation are publicly available on Zenodo (Version 1.0) at https://zenodo.org/records/23042604 (DOI: 10.5281/zenodo.23042604).

## Author contributions statement

**Ma.P**. and **S.A**. contributed equally to this work and jointly conceived the original research idea. **Ma.P**. developed the methodology, implemented the computational framework, and drafted the manuscript. **S.A**. collected and curated the data and participated in manuscript preparation and revision. **K.C**. contributed to software implementation and conducted the computational experiments. **Mo.P**. designed and prepared the figures and visualizations. **D.X**. supervised the project, provided scientific guidance and direction throughout the study, contributed to the interpretation of the results, and reviewed and revised the manuscript. All authors read and approved the final manuscript.

## Key Points

- We establish a ligand-aware, three-regime benchmark for residue-level protein–ligand binding-site prediction spanning *overrepresented, underrepresented*, and *zero-shot* ligand evaluation regimes.
- We use a progressive three-stage modeling suite that systematically evaluates the transition from single-ligand to multi-ligand and unseen-ligand residue-level prediction using protein sequences and ligand SMILES.
- In the original comparison, Stage 2 achieved higher overrepresented-test ligand-macro *F*_1_ than Stage 1 (0.5832 versus 0.4769), with simultaneous changes in predictor architecture, ligand conditioning, and the training regime.
- Ligand-input controls using constant, mismatched, and scrambled SMILES consistently reduce Stage 3 performance across all three evaluation regimes, supporting a contribution from the supplied ligand information rather than reliance solely on protein-derived or generic ligandability signals.
- Increasing protein language model scale generally improves performance across the evaluated configurations, although the trend is not strictly monotonic, whereas increasing chemical language model scale provides smaller and less consistent gains.

## Acknowledgments

This work was partially supported by the National Institutes of Health (grant R35GM126985).

## Abbreviations

CLM: Chemical Language Model
PLM: Protein Language Model
MLM: Masked Language Model
MLP: Multilayer Perceptron
CNN: Convolutional Neural Network
GNN: Graph Neural Network
ESM-2: Evolutionary Scale Modeling 2

## Extended Related Work

In this section, we categorize existing protein–ligand binding site prediction studies into distinct groups. For each study, we briefly describe two key aspects: the structure of the proposed method and the strategy used to address the class imbalance between binding and non-binding sites, which remains one of the main challenges in developing effective deep learning models for this task. While some previous works have grouped prediction methods into sequence- and structure-based approaches Roche et al. [2015], Xia et al. [2024], our focus is primarily on sequence-based methods. We further classify these studies into three groups based on the techniques used and the order in which they were developed. The earliest approaches applied traditional machine learning methods Yu et al. [2013], Yang et al. [2013], followed by deep learning-based methods Cui et al. [2019], Xia et al. [2020], and more recently, pretrained model-based methods Yuan et al. [2022], Zhang and Xie [2023]. We also include some studies that combine sequence- and structure-based methods due to their novelty in extracting sequence features or addressing the class imbalance problem, as these provide valuable insights and broaden the scope for the reader.

### Traditional Machine Learning Methods

Yu *et al*. Yu et al. [2013] proposed TargetS, which integrates multiple features for binding residue prediction, including position-specific scoring matrices (PSSMs) to capture protein evolutionary information, predicted secondary structures obtained using PSIPRED, and ligand-specific binding propensities. The ligand-specific propensities are calculated based on the frequencies of amino acids among binding sites for each ligand type. A 17-dimensional window centered at the target residue is used to capture not only the residue’s own propensity but also the influence of its local environment. To enhance prediction accuracy, the authors employed a modified AdaBoost (MAdaBoost) ensemble scheme that combines multiple classifiers. To address the class imbalance between binding and non-binding sites, they applied a random undersampling strategy within the AdaBoost framework.

Yang *et al*. Yang et al. [2013] proposed TM-SITE and S-SITE, two template-based methods that leverage structural and sequence similarity, respectively, to predict ligand-binding sites. TM-SITE performs structural comparisons by aligning the subsequence spanning from the first to the last binding residue (SSFL) of the query protein against a pre-calculated SSFL library using TM-align. Proteins with the highest alignment scores are selected as putative templates. A secondary scan is then conducted using the entire query structure against the BioLiP database. Final predictions are obtained through a clustering and voting scheme. In contrast, S-SITE relies on sequence-based alignment. PSI-BLAST is used to generate multiple sequence alignments for the query protein, from which a position-specific frequency matrix (PSFM) is derived. Template profiles in the BioLiP library, represented as position-specific scoring matrices (PSSMs), are precomputed in advance. The PSFM of the query is then compared to these template PSSMs using sequence alignment techniques, and a voting scheme is applied to determine the final binding sites.

Hu *et al*. Hu et al. [2016] introduced two ligand-specific methods for predicting ion-binding sites. IonSeq is a sequence-based approach that employs sequence profiles together with a modified AdaBoost algorithm to distinguish between binding and non-binding sites. IonCom is a composite method that integrates IonSeq with multiple template-based predictors, including COFACTOR, TM-SITE, S-SITE, and COACH, to improve prediction accuracy and robustness. To address class imbalance and reduce the risk of overfitting, both methods use a modified AdaBoost framework that applies selective random sampling to negative (non-binding) residues while fully utilizing all positive (binding) residues in every training round.

Hu *et al*. Hu et al. [2018] proposed ATPbind, a method that combines sequence-based features—including position-specific scoring matrices (PSSMs), predicted secondary structure, and solvent accessibility—with the outputs of S-SITE and TM-SITE, which are sequence- and structure-template-based predictors. To address the imbalance between binding and non-binding sites, ATPbind employs multiple support vector machines (SVMs) trained with a random undersampling strategy and integrates their outputs using a mean-ensemble approach.

Lin *et al*. Lin et al. [2016] presented the MIB webserver, which utilizes available templates from the Protein Data Bank (PDB) to identify binding sites for 12 types of metal ions. The method compares the structure of query proteins with these templates without any model training. Metal ion–binding residue templates are extracted from the PDB, and homologous proteins are filtered out to ensure diversity. The server aligns the query protein against the template structures and assigns a binding score based on sequence and structural similarity. MIB also predicts the docking positions of metal ions in protein structures.

### Deep Learning-Based Methods

Cui *et al*. Cui et al. [2019] introduced DeepCSeqSite (DCS-SI), a novel sequence-based approach for predicting protein–ligand binding sites. DCS-SI is built on a deep convolutional neural network with an encoder–decoder architecture. The encoder transforms entire amino acid sequences into hierarchical representations that capture both local and long-range dependencies between residues. The decoder then uses these features to predict binding sites. To address the class imbalance issue, the authors did not apply explicit oversampling or undersampling techniques. Instead, they leveraged the strong representation capability of their deep convolutional network and processed entire amino acid sequences as input units during mini-batch grouping. This strategy preserved the natural proportion of positive (binding) and negative (non-binding) samples within each mini-batch, reducing the risk of batches dominated by negative samples and enabling the model to learn from a more representative distribution.

Xia *et al*. Xia et al. [2020] proposed DELIA, a hybrid deep learning model that combines 1D sequence-based features with a 2D structure-based distance matrix representing the Euclidean distances between residues in a protein structure. The model is composed of three main components: a feature extractor, a ResNet module, and a BiLSTM module. The feature extractor utilizes tools such as PSI-BLAST, HHblits, SCRATCH-1D, and S-SITE to generate diverse sequence and profile features, which are then concatenated. The ResNet and BiLSTM modules are used to extract high-level representations, and their outputs are concatenated and passed through a softmax layer to generate the final predictions. To address the class imbalance issue, the authors applied a combination of random undersampling and oversampling. For undersampling, they constructed multiple training subsets, each containing all positive samples and 20% of the original negative samples. For oversampling, they ensured that each negative sample was used only once per epoch, while positive samples were randomly selected and reused multiple times across mini-batches.

Essien *et al*. Essien et al. [2019] utilized a Capsule Network (Caps-Net) for predicting zinc-binding sites. The model processes protein sequences using a sliding-window approach, where each window consists of 25 amino acids. These fixed-length segments are extracted from longer sequences and processed individually. Convolutional layers extract increasingly abstract features, while the PrimaryCaps layer captures higher-level representations. Dynamic routing further refines these features, and the output layer uses them to predict zinc-binding sites. To address class imbalance, the authors applied a bootstrapping strategy: in each iteration, a classifier was trained on a balanced subset containing all positive samples and a portion of negative samples.

This process was repeated multiple times so that all negative samples were eventually utilized, and the final prediction was obtained by averaging across the independently trained classifiers.

Xia *et al*. Xia et al. [2021] proposed GraphBind, a model that leverages both structure-based and sequence-based features to identify binding sites. GraphBind consists of three main modules: feature extraction, structural context extraction, and graph construction. Sequence features include position-specific scoring matrices (PSSMs) and hidden Markov models (HMMs), generated using PSI-BLAST and HHblits. Structure features include atomic properties and secondary structure information derived from DSSP. The structural context extraction module defines a local 3D environment around each target residue using a sliding sphere, producing pseudo-positions that represent nearby spatial regions. In the graph construction module, each residue is treated as a node, and a graph is built using node feature vectors, distance matrices, adjacency matrices, and edge features. This graph is then processed by a classifier to determine whether each residue is part of a binding site. To address class imbalance, sequence identity and structural similarity were first assessed with BL2SEQ and TM-align; chains with identity *>* 0.8 and TM-score *>* 0.5 were clustered, and binding annotations were transferred to a representative chain. Redundancy was further reduced with `CD-HIT` to ensure training-set sequence identity below 30%.

Xia *et al*. Xia et al. [2022] presented BindWeb, a model that integrates predictions from GraphBind and DELIA using mean-shift clustering to identify binding sites. Binding scores from GraphBind and DELIA are averaged to generate final predictions. For each residue, the averaged score is compared to an averaged threshold (based on the individual optimal thresholds of both methods) to determine whether the residue is binding or non-binding. Because BindWeb incorporates DELIA, it inherits DELIA’s random undersampling–based ensemble strategy for addressing class imbalance.

Essien *et al*. Essien et al. [2025] proposed GPred, a model with four modules for predicting coordinated metal ion–ligand binding sites. The *Point Neighborhood Grouper* identifies nearby atoms around each atom in a protein’s point cloud. The *Point Transformer* integrates atomic properties, geometric shape information, and evolutionary profiles using position encoding and self-attention. The *Residual Pooler* aggregates atomic-level features into residue-level representations through MaxPooling. Finally, a classifier predicts whether each residue is a binding site. To mitigate class imbalance, the authors used a weighted binary cross-entropy loss function.

Chelur *et al*. Chelur and Priyakumar [2022] proposed BiRDS, a deep residual neural network for predicting a protein’s most active binding site using sequence information alone. The model extracts a rich set of features, including token embeddings, positional embeddings, segment embeddings, position-specific scoring matrices, information content, secondary structure, and solvent accessibility. These features are concatenated and passed through the BiRDS network for classification. To address the imbalance between binding and non-binding residues, the authors trained the model using a weighted binary cross-entropy loss.

### Pretrained Model-Based Methods

Yuan *et al*. Yuan et al. [2022] proposed LMetalSite, a method in which the protein sequence is processed by a pretrained protein language model to generate embedding representations. To reduce overfitting, Gaussian noise is added to these embeddings during training. The noise-augmented embeddings are passed through shared transformer networks composed of multiple layers, after which ion-specific multi-layer perceptrons (MLPs) predict the binding patterns of four metal ions. To mitigate limited training data and exploit shared patterns among ions, the model adopts a multi-task learning framework.

Essien *et al*. Essien et al. [2023] proposed IonPred, a two-part framework for ion-binding site prediction. The first component is a masked language model (MLM) generator that predicts masked amino acids based on their sequence context. The second component is an ELECTRA-based discriminator that takes the (generator-modified) full sequence and determines whether each residue is original or replaced. To construct training samples, the authors computed amino acid frequency distributions for each ligand type to identify candidate residues and used sliding windows centered on these candidates to define positive and negative samples.

Pourmirzaei *et al*. Pourmirzaei et al. [2025a] proposed Prot2Token, a model that combines a protein language model with an autoregressive transformer. Protein sequences and task tokens (representing ligand types) are provided as input. The protein encoder processes the sequence, and its output—combined with positional embeddings—is projected and used by the decoder along with the task-token embeddings. The decoder is autoregressive and is pre-trained in a self-supervised manner, an important component when training data are limited.

Littmann *et al*. Littmann et al. [2021] proposed bindEmbed21, which consists of two modules: bindEmbed21DL (deep learning) and bindEmbed21HBI (homology-based inference). bindEmbed21DL uses embeddings from the Transformer-based ProtT5 model, which are fed into a shallow two-layer convolutional neural network. bindEmbed21HBI leverages the observation that proteins with high sequence similarity often share similar binding sites. The final bindEmbed21 model combines both modules. To address class imbalance, the authors used a weighted cross-entropy loss with ligand-specific weights.

Wang *et al*. Wang et al. [2024] introduced CLAPE-SMB, a model designed to predict generic small-molecule binding sites rather than ligand-specific sites. ESM-2 is used to extract protein features, and its parameters remain fixed during training to reduce computational cost and avoid catastrophic forgetting. The classifier is a five-layer multilayer perceptron (MLP) that predicts binding or non-binding residues. To mitigate class imbalance, the authors employed a composite loss combining class-balanced focal loss and triplet center loss.

Zhang *et al*. Zhang and Xie [2023] proposed LaMPSite, which takes both a protein’s amino acid sequence and a ligand’s 2D molecular graph as input. The protein sequence is processed by ESM-2 to generate residue embeddings and an estimated contact map. The ligand graph is processed using a graph neural network (GNN) to obtain atom embeddings and a distance map from a fast conformer. Each residue is paired with each atom to create an interaction map, which is refined using both the protein contact map and ligand distance map. Interaction scores for each residue are then averaged over ligand atoms, and high-scoring residues are grouped into predicted binding pockets using the protein contact map.

## Data Pre-Processing Details

BioLiP2 Zhang et al. [2024] was utilized as a comprehensive and curated database of biologically relevant protein–ligand binding interactions. The database primarily sources data from the Protein Data Bank (PDB), supplemented with annotations from curated literature and specialized databases such as Binding MOAD Benson et al. [2007] and BindingDB Liu et al. [2007]. Three main files were used in this work. The file BioLiP nr.txt.gz provides annotations for each ligand–protein interaction site, including binding residues in both original PDB numbering and renumbered indices starting from 1, along with metadata such as structure resolution and binding affinity. The file protein nr.fasta.gz contains receptor protein sequences clustered at 90% identity, formatted in FASTA, where header lines begin with “*>*” (e.g., *>*1hv2A) followed by amino acid sequences. The file ligand.tsv.gz contains ligand-level information, including Chemical Component Dictionary (CCD) identifiers, chemical formulas, SMILES strings, and other ligand descriptors. These resources were used to retrieve protein sequences, ligand SMILES strings, and residue-level binding annotations.

Sequences shorter than 50 residues were removed. Ligands lacking SMILES strings were also excluded. Binary residue-level labels were generated using the `binding_residues_renum` column: all residue positions were initialized to zero (non-binding), and positions corresponding to annotated binding residues were set to one (binding).

Ligands were then categorized by sample count:

- **Overrepresented:** ligands with at least 100 samples,
- **Underrepresented:** ligands with 20–99 samples,
- **Zero-shot:** ligands with 1–19 samples.

For each ligand, a separate FASTA file was generated containing all sequences that bind to that ligand. Each FASTA entry consisted of the sequence identifier and its amino acid sequence.

To reduce redundancy, `CD-HIT` was applied separately to each ligand-specific FASTA file using a 40% sequence identity threshold. CD-HIT clustering was performed independently within each ligand-specific receptor set to reduce within-ligand sequence redundancy.

Clusters were then assigned to training, validation, and test sets according to the predefined split ratios for each ligand category. After splitting, the *overrepresented* and *underrepresented* training sets were merged into a single unified training set, and their validation sets were combined into one shared validation set. All *zero-shot* ligands and their associated samples were reserved exclusively for testing and were not included in either training or validation. For evaluation, we retained three separate test sets: the *overrepresented, underrepresented*, and *zero-shot* test sets. This per-ligand clustering procedure reduced sequence redundancy within individual ligand-specific receptor datasets but did not enforce global receptor-sequence independence across different ligands or data splits.

The Chemical Component Dictionary (CCD) was used to retrieve SMILES strings for each ligand from `ligand.tsv.gz.` This file contains multiple SMILES variants, although the BioLiP2 documentation does not specify what each variant represents. For consistency, we used the first SMILES string provided. A comparison with RCSB PDB Berman et al. [2000] confirmed that this first entry corresponds to the standard SMILES and Canonical SMILES generated by OpenEye OEToolkits.

The resulting dataset contained 5,780 ligands and 41,327 samples. For systematic evaluation, ligands were categorized according to their sample counts. *Overrepresented* ligands were divided into two subgroups:

1. Ligands with at least 200 samples: this subgroup contained 32 ligands, with target training, validation, and test proportions of 70%, 10%, and 20%, respectively.
2. Ligands with 100–199 samples: this subgroup contained 13 ligands, with target training, validation, and test proportions of 50%, 20%, and 30%, respectively.

*Underrepresented* ligands had 20–99 samples. This category initially contained 123 ligands, with target training, validation, and test proportions of 34%, 33%, and 33%, respectively.

`CD-HIT` was applied separately within each ligand-specific receptor dataset at a 40% sequence-identity threshold to reduce sequence redundancy. The resulting clusters were then assigned to the training, validation, and testing sets according to the predefined splitting ratios. Because clustering was performed independently for each ligand, this procedure did not enforce global receptor-sequence independence across different ligands or evaluation groups. Thus, receptor sequences or related receptor clusters could occur in different splits when associated with different ligands.

After clustering, the next step is to assign clusters to the training, validation, and testing sets according to the predefined splitting ratios. However, for ligands with a small number of highly similar sequences, `CD-HIT` may produce fewer than three clusters. In such cases, there are not enough clusters to split across training, validation, and testing sets properly. This occurred for only one ligand in the *underrepresented* category (`CL0`), which is why it was excluded. For the *zero-shot* category, `CD-HIT` clustering was not used for dataset partitioning because all samples in this category were reserved exclusively for testing.

The original dataset split statistics after exclusion of CL0 are summarized in Table 7. These describe the original dataset version, distinct from the released modified Stage 3 dataset summarized in Table 1.

**Table 7.**
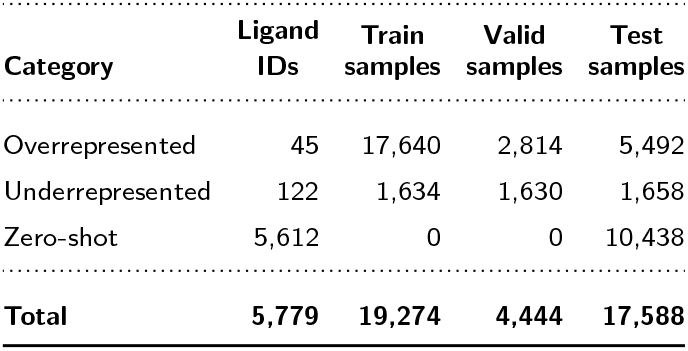
Original dataset split statistics after exclusion of CL0, before SMILES compatibility modifications. Counts reproduce the original dataset statistics.

The third category, *zero-shot*, comprised 5 612 ligands with 1–19 samples each. All associated samples were reserved exclusively for testing; none were included in training or validation.

During integration with the UniMol-2 CLMs, we applied minor changes to the dataset to ensure compatibility. UniMol-2’s built-in SMILES parser imposed stricter parsing constraints than the MolFormer tokenizer/parser. To standardize ligand representation in different CLMs, we attempted to use a standard canonical SMILES generated by OpenEye OEToolkits Hawkins et al. [2007]. However, some SMILES representations under this format were rejected by the UniMol-2 parser. In such cases, we attempted to substitute alternative SMILES variants. These alternatives were selected from the additional SMILES strings listed in the `ligand.tsv.gz file`. If no alternatives were accepted or available, the ligand was excluded from the dataset. The ligand `TRP`, which appears in the *underrepresented* datasets, required SMILES modification to be parsed successfully. In the *zero-shot* test set, 88 ligand IDs were affected by the SMILES compatibility procedure: 59 were retained with changed SMILES representations, while 29 had no valid alternatives and were excluded from the final *zero-shot* test set.

The excluded ligands from the *zero-shot* test set included `08T, 7RZ, 8M0, AOH, B1M, COB, D6N, DVT, GCR, ICS, K6G, KEG, MO7, NFV, OEX, OZN, PQJ, R5Q, S5Q, S9F, SIW, TEW, U00, UZC, V9G, WJS, WO2, X3P`, and `ZRW`.

### Ligand Composition and Molecular-Size Summary

To further characterize the chemical composition of the modified Stage 3 evaluation datasets, we summarized ligand type, molecular size, and chemical input length across the overrepresented, underrepresented, and zero-shot test groups. These summaries were calculated using the exact modified Stage 3 test datasets used for evaluation.

Ligands were assigned to four mutually exclusive chemical classes: metal or single-atom ions, other inorganic species, organic or carbon-containing molecules, and unclassified ligands. Classification was based on the evaluated SMILES representations together with current Chemical Component Dictionary metadata. The organic or carbon-containing category is a broad operational class that includes larger and organometallic ligands and should not be interpreted as a drug-likeness classification or molecular-size cutoff.

As shown in Table 8, organic or carbon-containing ligands constitute the majority of ligand IDs in all three groups, increasing from 73.33% of the overrepresented group to 86.78% of the underrepresented group and 97.13% of the zero-shot group. The zero-shot group also contains substantially more ligand IDs than the represented groups.

**Table 8.** Chemical composition of the modified Stage 3 test datasets. Percentages are calculated using ligand IDs within each test group rather than protein–ligand sample rows. The chemical classes are mutually exclusive. Organic or carbon-containing is a broad operational category that includes larger and organometallic ligands. The 15 unclassified zero-shot ligand IDs account for 23 sample rows and remain in the evaluation.

| **Ligand class** | **Overrepresented** | **Underrepresented** | **Zero-shot** |
|---|---|---|---|
| Metal or single-atom ion | 8 (17.78%) | 4 (3.31%) | 34 (0.61%) |
| Other inorganic | 4 (8.89%) | 12 (9.92%) | 111 (1.99%) |
| Organic or carbon-containing | 33 (73.33%) | 105 (86.78%) | 5,423 (97.13%) |
| Unclassified | 0 (0.00%) | 0 (0.00%) | 15 (0.27%) |
| Total ligand IDs | 45 | 121 | 5,583 |
| Protein–ligand sample rows | 5,527 | 1,809 | 10,365 |

**Table 9.**
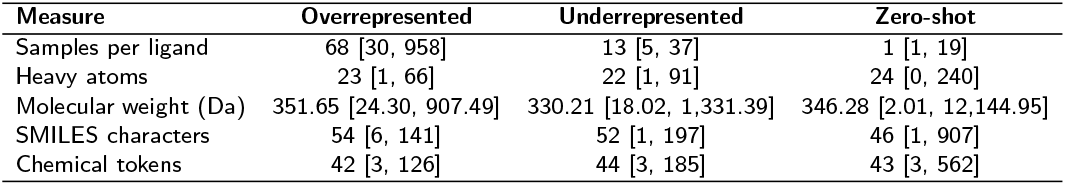
Ligand size and chemical-input-length summary for the modified Stage 3 test datasets. Values are median [minimum, maximum] across ligand IDs and are not sample-weighted summaries or interquartile ranges. Heavy-atom counts and molecular weights were calculated from the evaluated SMILES. Chemical-token counts include special tokens before truncation.

| **Measure** | **Overrepresented** | **Underrepresented** | **Zero-shot** |
|---|---|---|---|
| Samples per ligand | 68 [30, 958] | 13 [5, 37] | 1 [1, 19] |
| Heavy atoms | 23 [1, 66] | 22 [1, 91] | 24 [0, 240] |
| Molecular weight (Da) | 351.65 [24.30, 907.49] | 330.21 [18.02, 1,331.39] | 346.28 [2.01, 12,144.95] |
| SMILES characters | 54 [6, 141] | 52 [1, 197] | 46 [1, 907] |
| Chemical tokens | 42 [3, 126] | 44 [3, 185] | 43 [3, 562] |

The zero-shot group contains substantially fewer samples per ligand than the overrepresented and underrepresented groups. In contrast, median heavy-atom counts and molecular weights are broadly similar across the three groups, although their observed ranges differ substantially. These statistics describe differences among the evaluation groups but do not, by themselves, establish the cause of differences in predictive performance.

## Receptor Sequence Diversity Analysis

To characterize receptor-sequence diversity beyond the ligand-frequency definitions used to construct the evaluation regimes, we performed a receptor-diversity analysis using the exact modified Stage 3 test datasets used for the reference evaluation. These datasets contain 17,701 protein–ligand sample rows and 5,749 ligand IDs across the overrepresented, underrepresented, and zero-shot test groups, as summarized in Table 10.

**Table 10.** Receptor-sequence diversity summary for the exact modified Stage 3 test datasets. Distinct receptor-sequence counts are calculated within each evaluation group. Local CD-HIT clusters were generated separately for each ligand at 40% sequence identity; therefore, the summed cluster counts do not represent globally unique receptor families.

| Test Group | Sample Rows | Distinct Receptor Sequences | Ligand IDs | Sum of Local Clusters |
|---|---|---|---|---|
| Overrepresented | 5,527 | 5,352 | 45 | 3,398 |
| Underrepresented | 1,809 | 1,719 | 121 | 1,305 |
| Zero-shot | 10,365 | 9,550 | 5,583 | 9,114 |

For each ligand, distinct full receptor sequences were identified and clustered separately using CD-HIT v4.8.1 at 40% sequence identity (`-c` 0.4), with word length 2 (`-n` 2) and global identity (`-G` 1). Sequences were ordered by decreasing full sequence length and then by sequence hash. Exact duplicate receptor sequences were collapsed only for the clustering analysis; all original sample rows were retained in the performance evaluation. Because the historical clustering files used during the original dataset construction are unavailable, these clusters represent a fresh analysis of the frozen evaluation datasets rather than a reconstruction of the original split assignments.

For each ligand, normalized receptor diversity was defined as

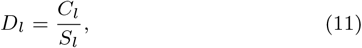

where *C*_*l*_ is the number of CD-HIT clusters and *S*_*l*_ is the number of distinct full receptor sequences associated with ligand *l*. Thus, lower values indicate greater sequence redundancy among the receptors associated with a ligand, whereas values closer to 1 indicate that a larger fraction of the distinct receptor sequences form separate clusters. This metric measures receptor-sequence redundancy and should not be interpreted as a direct measure of structural diversity, binding-site diversity, or prediction difficulty.

### Treatment of singleton ligands

Ligands associated with only one distinct receptor sequence require special treatment when interpreting normalized receptor diversity. For such a ligand, *C*_*l*_ = *S*_*l*_ = 1, and therefore *D*_*l*_ = 1 by definition. Consequently, a diversity value of 1 for a singleton does not provide evidence that the ligand is associated with a diverse set of receptors.

As summarized in Table 12, this issue is particularly important for the zero-shot group. All overrepresented and underrepresented ligands have more than one distinct receptor sequence, whereas 4,001 of the 5,583 zero-shot ligand IDs (71.7%) are singletons. The remaining 1,582 zero-shot ligands have at least two distinct receptor sequences. We therefore retain all singleton ligands in the dataset summaries and performance analyses, but report them separately when characterizing receptor-diversity distributions and exclude them from the diversity boxplot. This prevents the zero-shot distribution from being dominated by values of 1 that arise mechanically from having only one observed receptor sequence.

**Table 11.** Zero-shot performance for singleton and non-singleton ligands. Singletons have one distinct full receptor sequence; non-singletons have more than one. Results use saved predictions from one released Stage 3 checkpoint. All ligand IDs and sample rows, including zero-*F*_1_ ligands, are retained. The comparison is descriptive, and subgroup uncertainty intervals were not estimated.

| Subgroup | Ligand IDs | Samples | Pooled *F*<sub>1</sub> | Ligand-macro *F*<sub>1</sub> |
|---|---|---|---|---|
| Singleton | 4,001 | 4,009 | 0.2964 | 0.2312 |
| Non-singleton | 1,582 | 6,356 | 0.3336 | 0.2548 |

**Table 12.** Per-ligand receptor-diversity summary. Singleton ligands have only one distinct receptor sequence and are reported separately because their normalized diversity is necessarily 1. Median diversity and interquartile ranges are therefore calculated for non-singleton ligands.

| Test Group | Non-Singleton Ligands | Singleton Ligands | Median Diversity [Q1, Q3] |
|---|---|---|---|
| Overrepresented | 45 | 0 | 0.639 [0.570, 0.718] |
| Underrepresented | 121 | 0 | 0.769 [0.600, 0.875] |
| Zero-shot | 1,582 | 4,001 | 1.000 [0.667, 1.000] |

Single-checkpoint pooled *F*_1_ was 0.2964 for singleton ligands and 0.3336 for non-singletons. Ligand-macro *F*_1_ was 0.2312 and 0.2548, respectively (Table 11). We report the groups separately because a per-ligand score based on one receptor has limited support. The subgroup comparison is descriptive.

A singleton denotes one distinct full receptor sequence, rather than one dataset row or one CD-HIT cluster. Singleton ligands accounted for 4,009 sample rows because a ligand with only one distinct receptor sequence could occur in multiple rows. All 5,583 ligand IDs and 10,365 samples were retained. Pooled *F*_1_ combines valid-residue TP, FP, and FN counts, whereas ligand-macro *F*_1_ gives each ligand ID equal weight after pooling its own residue counts. Zero-*F*_1_ ligands were retained. Scores use saved predictions from one released Stage 3 checkpoint, a probability threshold strictly greater than 0.5, and valid residues within the first 1,024 residues. The groups contain different proteins and ligands, and no subgroup uncertainty intervals were estimated. These results do not establish that receptor count causes the performance difference.

The three evaluation regimes exhibited different observed receptordiversity distributions among ligands with multiple distinct receptor sequences (Figure 6). Median normalized diversity was 0.639 for the overrepresented group, 0.769 for the underrepresented group, and 1.000 for the non-singleton zero-shot group. However, these values should be interpreted together with receptor counts and singleton prevalence. In particular, 71.7% of zero-shot ligand IDs were associated with only one distinct receptor sequence and therefore cannot provide an informative within-ligand estimate of receptor diversity.

**Figure 6.**
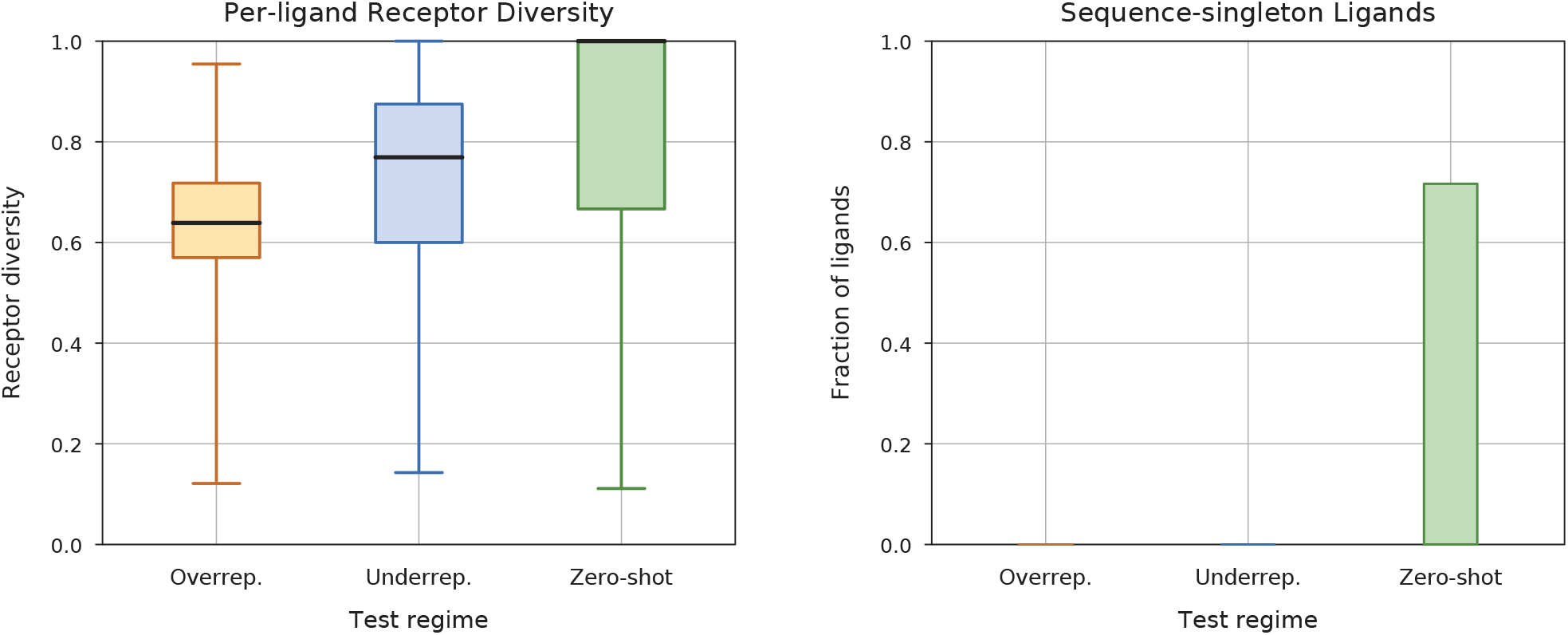
Receptor sequence diversity in the modified Stage 3 test datasets. The left panel shows normalized receptor diversity for ligands with at least two distinct receptor sequences; boxes indicate the interquartile range, central lines indicate the median, and whiskers span the minimum and maximum. The right panel shows the fraction of singleton ligands in each evaluation group. All 4,001 zero-shot singleton ligands (71.7% of 5,583 zero-shot ligand IDs) are retained in the dataset summaries and performance analyses but are excluded from the diversity boxplot because their normalized diversity is necessarily 1.

Accordingly, the overrepresented, underrepresented, and zero-shot regimes remain frequency-defined evaluation groups rather than categories defined by receptor diversity. The receptor-diversity analysis provides additional context regarding sequence redundancy within each regime; however, the normalized diversity ratio also depends on the number of associated receptor sequences, and a higher ratio does not by itself establish greater structural diversity, more complex binding behavior, or a more difficult prediction problem.

### Whole-receptor sequence diversity and performance

As complementary sequence-based evidence, we examined the relationship between whole-receptor sequence diversity and per-ligand prediction performance among non-singleton ligands (Fig. 7). Receptor diversity was defined as the number of local 40% CD-HIT clusters divided by the number of distinct full receptor sequences for each ligand. The Spearman correlations between receptor diversity and per-ligand F1 were −0.432, 0.092, and 0.041 for the overrepresented, underrepresented, and zero-shot groups, respectively. Thus, a moderate negative association was observed for the overrepresented group, whereas the associations were near zero for the underrepresented and zero-shot groups. These results describe whole-receptor sequence diversity and should not be interpreted as binding-site sequence conservation or as evidence that receptor diversity does not affect prediction performance.

**Figure 7.**
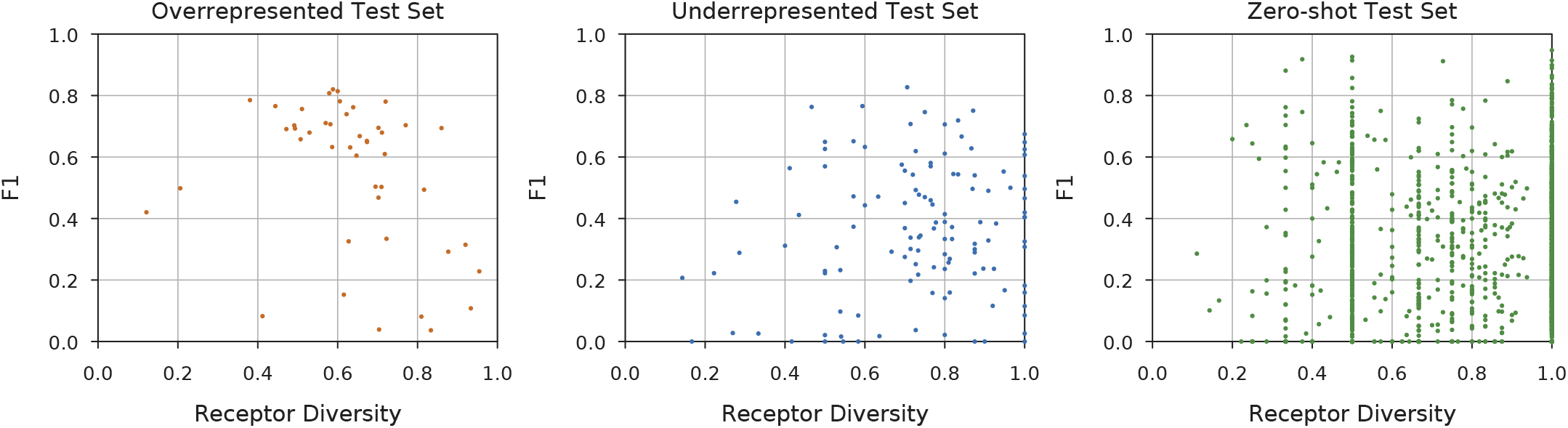
Full-receptor sequence diversity and ligand-level performance. Each point represents a non-singleton ligand (45 overrepresented, 121 underrepresented, and 1,582 zero-shot). Diversity is the local 40% CD-HIT cluster count divided by the number of distinct receptor sequences. F1 is pooled per ligand across its test rows. This analysis uses whole-receptor sequence diversity and is distinct from the binding-site sequence-identity metric and the structurally covered subset analyzed separately.

### Cross-split receptor overlap

Because the receptor-diversity analysis above clusters sequences separately for each ligand, it characterizes within-ligand redundancy but does not establish receptor independence across ligands or data splits. We therefore performed a separate cross-split audit using the union of the five frozen modified Stage 3 files: training, validation, and the overrepresented, underrepresented, and zero-shot test sets. The combined dataset contains 41,212 protein–ligand sample rows and 30,288 distinct full receptor sequences.

To provide a common receptor-family assignment across all ligand groups and splits, the 30,288 distinct receptor sequences were clustered jointly using CD-HIT at 40% sequence identity. This global analysis produced 15,925 clusters. These global clusters are distinct from the per-ligand clusters used for the receptor-diversity analysis above: per-ligand clustering quantifies redundancy among receptors associated with an individual ligand, whereas global clustering assigns a common analysis cluster ID across the union of all five files and therefore permits comparison of receptor co-membership across ligands and splits.

Three distinct forms of overlap should therefore be distinguished. First, ligand identity determines the evaluation regime: in the zeroshot setting, ligand IDs in the test set are absent from training and validation. Second, exact receptor-sequence overlap occurs when an identical full receptor sequence is present in both a test group and the supplied training dataset. Third, global-cluster overlap occurs when a test receptor and at least one training receptor are assigned to the same global CD-HIT cluster, even when their full sequences are not identical. Thus, zero-shot ligand identity does not imply that the corresponding receptor sequence or receptor cluster is unseen during training.

The audit shows that receptor reuse occurs across the released training and test files (Figure 8). Exact full-sequence overlap with training was observed for 23.60%, 35.89%, and 26.29% of distinct receptors in the overrepresented, underrepresented, and zero-shot test groups, respectively. When overlap was assessed using global CD-HIT cluster co-membership, the corresponding proportions were 48.90%, 56.43%, and 50.85%. Because same-cluster overlap includes exact sequence matches, these percentages represent nested rather than additive forms of overlap.

**Figure 8.**
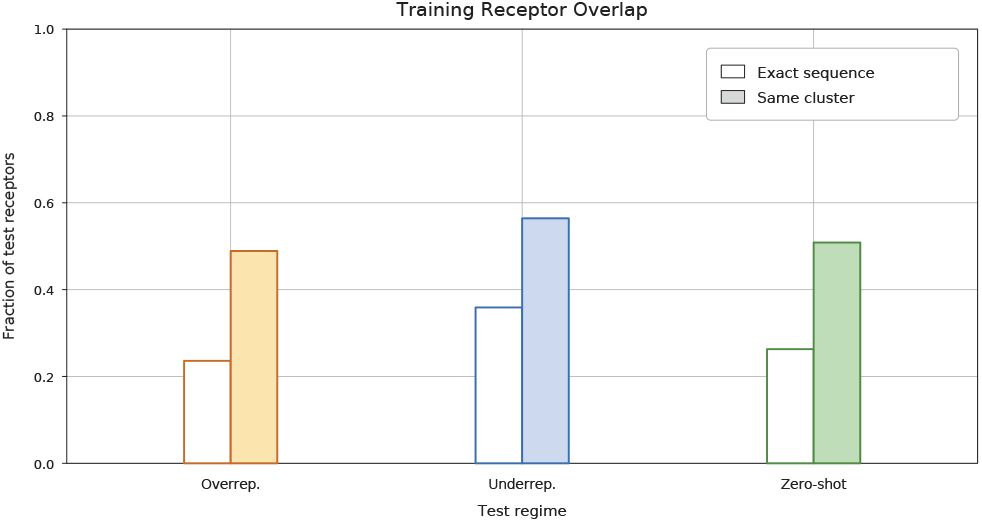
Exact and global-cluster receptor overlap with the supplied training dataset. Exact overlap denotes distinct test receptor sequences that also occur identically in the training dataset. Global-cluster overlap denotes distinct test receptor sequences assigned to a global 40% CD-HIT cluster containing at least one training receptor. The denominator is the number of distinct full receptor sequences in each test group. Same-cluster overlap includes exact sequence matches, so the two percentages are not additive.

We additionally compared zero-shot performance between receptor-overlapping and receptor-unseen subsets using saved Stage 3 predictions from one released checkpoint. For receptors with an exact sequence match in the released training set, pooled residue-level *F*_1_ was 0.2569, compared with 0.3470 for receptors without an exact match. Using the global 40% CD-HIT audit clusters, pooled residue-level *F*_1_ was 0.3140 for the overlapping subset (5,363 samples) and 0.3259 for the non-overlapping subset (5,002 samples).

Unlike receptor-cluster-balanced *F*_1_, this analysis directly compares performance according to training-receptor overlap. Performance did not decrease on the receptor-unseen subsets, suggesting that useful predictions are not restricted to receptors overlapping with the released training set. However, the subsets contain different proteins and ligands, and this single-checkpoint comparison is descriptive. It does not establish an absence of memorization. Here, receptor-unseen means no exact sequence match or no shared audit cluster with the released training set, depending on the criterion; it does not imply absence from protein-encoder pretraining.

These results clarify the scope of the evaluation. The zero-shot setting is defined at the ligand-identity level: zero-shot ligand IDs are excluded from training and validation. It is not a receptor-unseen or receptor-family-unseen benchmark. Likewise, per-ligand receptor clustering can reduce sequence redundancy within an individual ligand dataset but does not by itself enforce global receptor independence when the same receptor is associated with different ligands.

The global clusters used in this audit are fresh analysis clusters rather than recovered historical split-generation assignments, because the original cluster-assignment files are unavailable. CD-HIT cluster co-membership should also be interpreted as an operational sequence-similarity measure: membership in the same cluster does not guarantee that every pair of sequences within that cluster is directly at least 40% identical, while membership in different clusters does not prove the absence of cross-split homology. Accordingly, the observed receptor overlap documents properties of the released datasets but does not by itself establish label leakage, demonstrate model memorization, or reconstruct the exact historical checkpoint-training membership. No complete composite sample keys were shared across the five released files, where the composite key comprises PDB ID, receptor chain, row ID, and ligand ID. This record-level separation should not be interpreted as receptor-sequence separation, since different records can contain identical or globally clustered receptor sequences.

### Chemical similarity of zero-shot ligands

We compared the evaluated SMILES representations of all 5 583 zeroshot ligand IDs in the modified Stage 3 dataset against the 166 supervised-training ligand IDs. Canonical SMILES were generated using RDKit 2026.03.6. The primary representation-matching procedure retained encoded stereochemistry; no salt stripping, charge neutralization, or tautomer normalization was applied. There were 31 stereo-aware canonical representation matches. Of these, 30 contained unspecified stereochemistry, so these are matching input representations rather than confirmed identical stereoisomers. Ignoring stereochemistry identified 67 matches, including the 31 primary matches.

All performance results in this analysis use the same saved predictions from one released checkpoint, a probability threshold strictly greater than 0.5, and valid residues within the 1 024-residue evaluation window. Excluding the 31 primary matches changed pooled residue-level *F*_1_ from 0.3193 to 0.3179; excluding the 67 matches identified without stereochemistry changed it to 0.3166 (Table 13). These small changes indicate that matching input representations account for little of the aggregate performance. These single-checkpoint sensitivity results do not replace the three-seed benchmark results in Table 2.

**Table 13.**
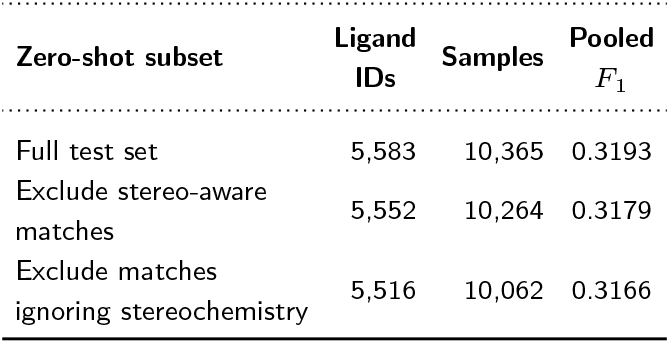
Chemical-overlap sensitivity analysis for the zero-shot test set using one released Stage 3 checkpoint. The stereochemistry-ignoring matches include the stereo-aware matches; the exclusions are applied separately.

For chemical similarity, we used chirality-aware Morgan fingerprints with radius 2 and 2 048 bits, calculating the maximum Tanimoto similarity to any supervised-training ligand. The similarity figures include 5 423 organic or carbon-containing zero-shot ligand IDs. Ions, other inorganic inputs, and unclassified IDs remain included in the full-set representation-matching and exclusion analyses but are omitted from these plots.

Median nearest-training similarity was 0.2405. The descriptive Spearman correlation between nearest-training similarity and per-ligand *F*_1_ was 0.1966, indicating a weak positive association. Pooled residue-level *F*_1_ increased from approximately 0.230 in the lowest-similarity bin to 0.443 in the highest. Ligand-macro *F*_1_ increased through the 0.6–0.8 bin and decreased slightly in the final bin (Figure 9). Per-ligand results are shown in Figure 10. Pooled residue-level *F*_1_ combines residue counts, whereas ligand-macro *F*_1_ gives each ligand ID equal weight. Because the bins contain different proteins and ligand compositions, these comparisons do not establish a causal effect of chemical similarity. They also do not demonstrate an absence of memorization or chemical-encoder pretraining exposure.

**Figure 9.**
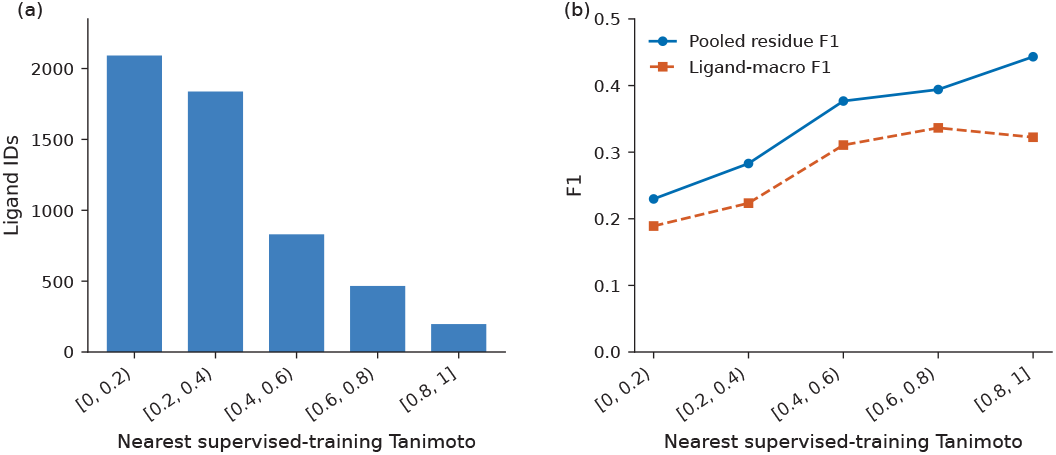
Chemical-similarity distribution and zero-shot performance. (a) Numbers of organic or carbon-containing zero-shot ligand IDs in five pre-defined nearest-training Tanimoto similarity bins. (b) Pooled residue-level *F*_1_ and ligand-macro *F*_1_ for the same bins. From left to right, bins contain 2,092, 1,838, 830, 466, and 197 ligand IDs and 2,775, 3,543, 1,864, 1,156, and 569 samples. The final bin includes similarity 1. Results use one released check-point. The comparison is descriptive and no confidence intervals are shown.

**Figure 10.**
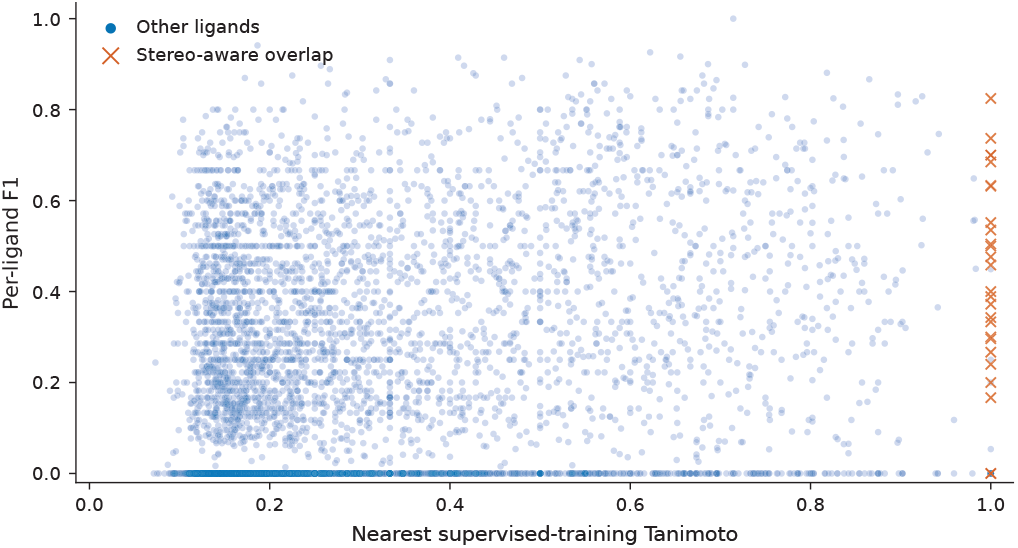
Chemical similarity and per-ligand zero-shot performance. Each point represents one of 5,423 organic or carbon-containing zero-shot ligand IDs. The horizontal axis shows the maximum chirality-aware Morgan Tanimoto similarity to the 166 supervised-training ligand inputs. Per-ligand *F*_1_ pools valid-residue counts across that ligand’s samples. Orange crosses mark the 31 stereo-aware canonical representation matches. All ligands, including those with zero *F*_1_ or only one sample, are retained. Similarity 1 alone does not establish chemical identity.

## Binding-Site Structural Variability Analysis

To characterize structural variability among experimentally observed binding sites associated with the same ligand, we performed a post-hoc analysis using available experimental reference structures for samples in the modified Stage 3 test datasets. This analysis was performed only for evaluation and did not provide structural information to the prediction model. Binding-site RMSD therefore measures variation among experimental reference binding sites associated with the same ligand, rather than structural error between model predictions and experimental structures. The analysis reused the saved Stage 3 predictions without model retraining, new inference, threshold modification, or reconstruction of the dataset splits.

Experimental X-ray and electron-microscopy structures with reported resolution no worse than 4 Å were considered, using the first structural model. Annotated binding residues were mapped through the receptor sequences and assigned to matching ligand instances using heavy-atom contacts within 5 Å. Ambiguous overlapping ligand-instance assignments were excluded from the primary structural comparisons.

Receptor-residue correspondence between structure pairs was established using US-align. Both length-normalized TM-scores were required to be at least 0.5. Binding sites were matched using mutual unique best correspondence overlap rather than by selecting the pair with the lowest RMSD. A valid comparison required at least four matched C*α* atoms and at least 50% coverage of each annotated binding site. Missing C*α* atoms remained in the coverage denominator. Comparisons with collinear fits and sites outside the saved first-1,024-residue evaluation window were excluded. Local binding-site RMSD was then calculated after a separate rigid-body superposition of the matched site C*α* atoms.

Each ligand contributed one observation to the structural-variability analysis. The ligand-level structural-diversity value was defined as the design-weighted median of the accepted receptor-pair median binding-site RMSDs. The weights accounted for sampling fractions in same- and different-receptor-sequence-cluster strata but did not correct for structural missingness. Per-ligand covered F1 was calculated by pooling the saved true-positive, false-positive, and false-negative counts over the unique sample rows contributing to accepted structural comparisons, so that a sample appearing in multiple receptor pairs was not counted repeatedly.

### RMSD and prediction performance

Valid ligand-level RMSD estimates were obtained for 36 overrepresented, 89 underrepresented, and 600 zero-shot ligands. We assessed the relationship between ligand-level binding-site RMSD and covered-subset *F*_1_ using Spearman correlation. Adjusted associations were calculated as partial Spearman correlations by rank residualization. We ranked binding-site RMSD, covered-subset *F*_1_, and each covariate using average ranks for ties. We separately regressed ranked RMSD and ranked *F*_1_ on the ranked covariates with an intercept using ordinary least squares. The Pearson correlation between the two residual vectors was reported as the partial Spearman correlation. The five covariates were log covered sample count, binding-site size, receptor sequence diversity, reported structure resolution, and training receptor-cluster overlap. Constant covariates were removed. Estimates required more than *k* + 10 ligands, where *k* is the number of retained covariates. These are descriptive adjustments, not causal estimates. Uncertainty intervals for the unadjusted Spearman correlations were estimated using 2,000 bootstrap draws with seed 20260917. Ligands sharing contributing PDB entries or global receptor clusters were connected into components, and whole components were resampled. The overrepresented, underrepresented, and zero-shot groups contained 28, 75, and 456 components, respectively. These exploratory intervals are conditional on the covered subset and are not corrected for multiple comparisons. The results are summarized in Table 14, and the ligand-level RMSD–*F*_1_ relationships are shown in Figure 11.

**Table 14.**
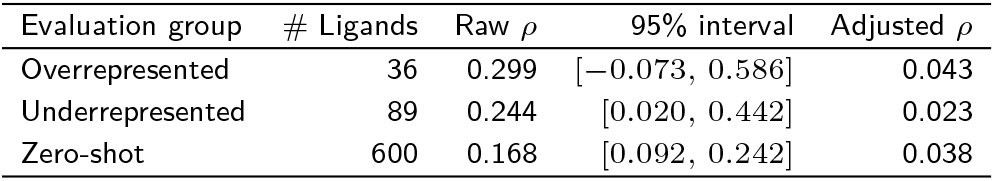
Association between experimental binding-site structural variability and prediction performance. Raw values are Spearman correlations between ligand-level binding-site RMSD and covered *F*_1_. Adjusted values are partial Spearman correlations obtained by rank residualization, accounting for log covered sample count, binding-site size, receptor-sequence diversity, reported structure resolution, and training-receptor cluster overlap. The 95% bootstrap intervals apply only to the raw correlations; no uncertainty intervals were computed for the adjusted values.

| Evaluation group | # Ligands | Raw ρ | 95% interval | Adjusted ρ |
|---|---|---|---|---|
| Overrepresented | 36 | 0.299 | [−0.073, 0.586] | 0.043 |
| Underrepresented | 89 | 0.244 | [0.020, 0.442] | 0.023 |
| Zero-shot | 600 | 0.168 | [0.092, 0.242] | 0.038 |

**Table 15.** Matched per-ligand comparison of the Stage 1 linear predictor and a Stage 2-compatible decoder on six selected overrepresented ligands. Values are mean *F*_1_± sample standard deviation across three paired random seeds. Differences are decoder minus linear, calculated within each seed before summarizing. Each ligand’s *F*_1_ pools residue counts within its test set. The six-ligand ligand-macro *F*_1_ is the unweighted mean of the six ligand scores, calculated separately for each seed. Evaluation includes padded positions in both ablation arms, unlike the main experiments. Differences are computed from unrounded results and therefore may differ slightly from subtraction of the displayed rounded means.

| Ligand | Linear<br>*F*<sub>1</sub> | Decoder<br>*F*<sub>1</sub> | Paired<br>difference |
|---|---|---|---|
| ZN | 0.8274 ± 0.0005 | 0.8285 ± 0.0023 | +0.0012 ± 0.0027 |
| ATP | 0.5871 ± 0.0070 | 0.5833 ± 0.0140 | −0.0038 ± 0.0071 |
| PLP | 0.8050 ± 0.0067 | 0.8291 ± 0.0043 | +0.0241 ± 0.0110 |
| GLC | 0.2613 ± 0.0148 | 0.3095 ± 0.0138 | +0.0483 ± 0.0285 |
| FE2 | 0.7442 ± 0.0281 | 0.7366 ± 0.0418 | −0.0076 ± 0.0386 |
| BMA | 0.0338 ± 0.0189 | 0.0532 ± 0.0922 | +0.0194 ± 0.1110 |
| **Six-ligand<br>Ligand-macro *F*<sub>1</sub>** | **0.5431 ± 0.0048** | **0.5567 ± 0.0198** | **+0.0136 ± 0.0171** |

**Table 16.** Paired bootstrap analysis of the ligand-input control experiments. Differences are calculated as native-SMILES pooled *F*_1_ minus control pooled *F*_1_. Confidence intervals were estimated using 2,000 paired bootstrap resamples of exact receptor-sequence groups within each test set.

| **Test group** | **Control** | **△ pooled *F*<sub>1</sub>** | **95% CI** |
|---|---|---|---|
| Overrepresented | Constant | 0.2969 | [0.2873, 0.3068] |
| Overrepresented | Mismatched | 0.3957 | [0.3855, 0.4063] |
| Overrepresented | Scrambled | 0.1222 | [0.1142, 0.1307] |
| Underrepresented | Constant | 0.0957 | [0.0843, 0.1061] |
| Underrepresented | Mismatched | 0.1539 | [0.1398, 0.1671] |
| Underrepresented | Scrambled | 0.1128 | [0.1002, 0.1254] |
| Zero-shot | Constant | 0.1858 | [0.1793, 0.1928] |
| Zero-shot | Mismatched | 0.0800 | [0.0739, 0.0863] |
| Zero-shot | Scrambled | 0.0609 | [0.0570, 0.0648] |

**Figure 11.**
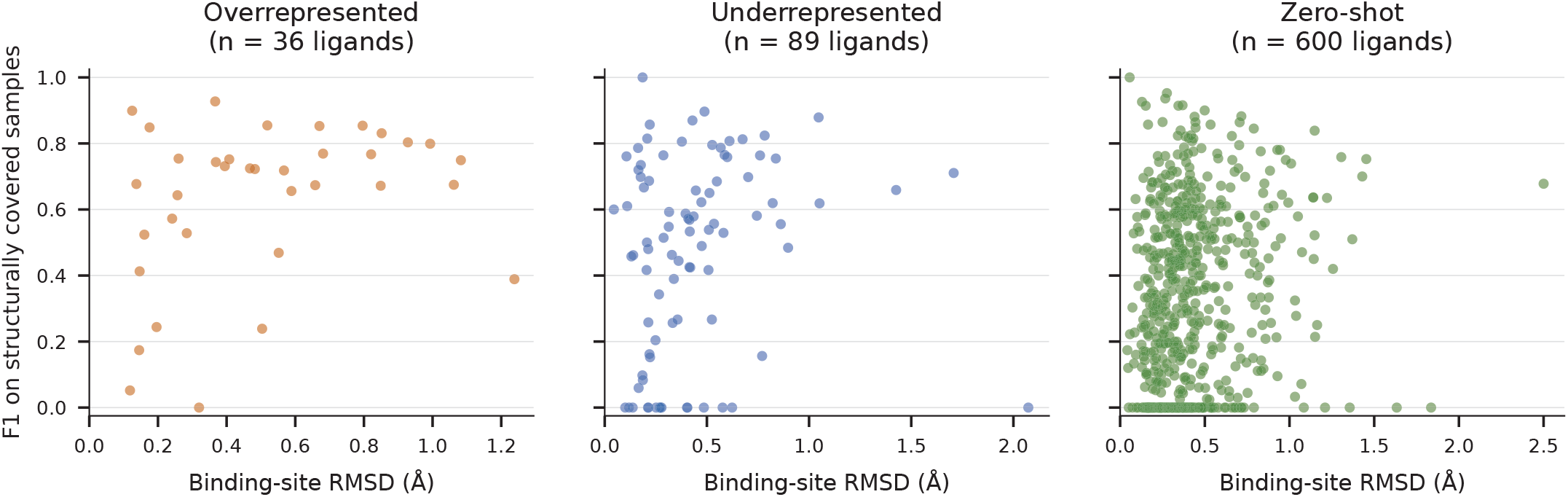
Relationship between experimental binding-site structural variability and prediction performance in the structurally covered subset. Each point represents one ligand. The x-axis shows the ligand-level binding-site RMSD derived from comparisons among available experimental reference binding sites associated with that ligand, and the y-axis shows the corresponding covered *F*_1_ calculated from samples contributing to accepted structural comparisons. RMSD quantifies variability among experimental reference binding sites and does not represent structural error between model predictions and experimental structures.

After adjustment, the correlations were 0.043, 0.023, and 0.038 for the overrepresented, underrepresented, and zero-shot groups, respectively. These descriptive results should be interpreted within the structurally covered subset and do not establish the absence of memorization. The unadjusted overrepresented-group estimate was based on only 36 ligands, and its wide 95% interval included zero, limiting the strength of conclusions for this group.

### Structural coverage and performance

The structural analysis was restricted to samples that passed the structural mapping and quality-control criteria. As shown in Fig. 12, the resulting structurally covered subsets represented only a portion of each test population and showed higher pooled residue-level *F*_1_ than the corresponding full test populations. Therefore, the structural analyses should be interpreted as characterizing the structurally eligible subset rather than the complete test populations.

**Figure 12.**
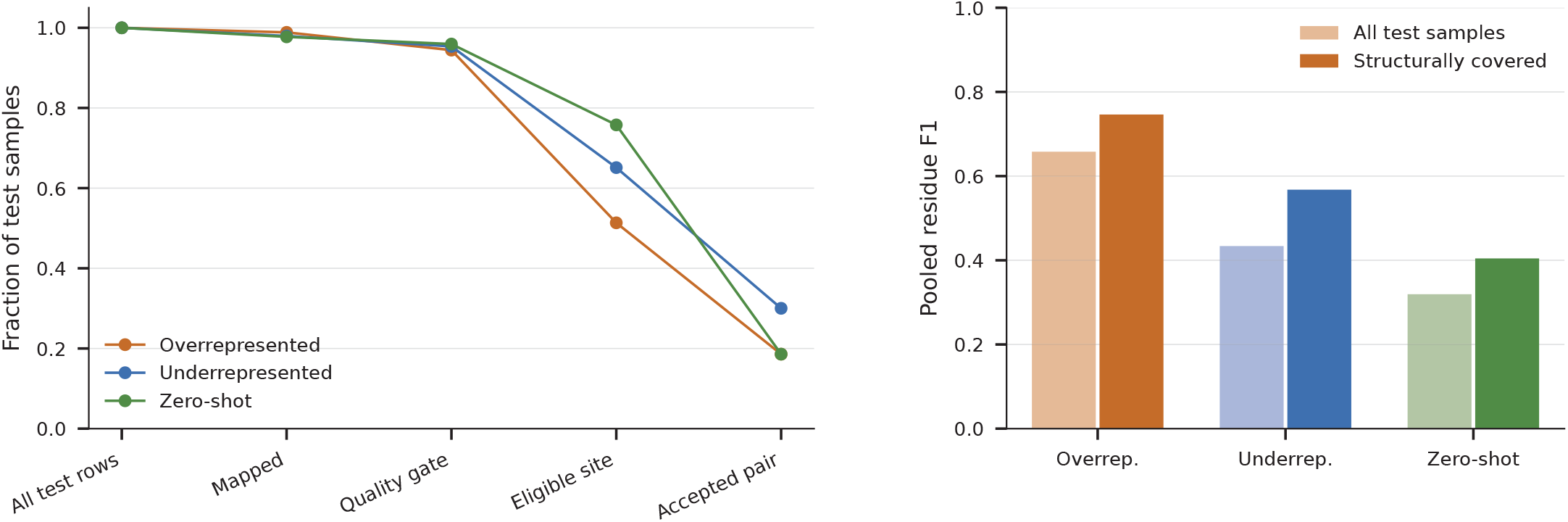
Structural coverage and performance. Left: fraction of test samples retained through successive stages of the structural mapping and quality-control pipeline for the overrepresented, underrepresented, and zero-shot groups. Right: pooled residue-level F1 for the full test populations compared with the structurally covered subsets. Structural coverage selected a higher-performing subset in all three evaluation groups.

### Binding-site sequence conservation

We additionally examined binding-site sequence conservation within the same structurally covered subset by relating reference-site amino-acid identity to per-ligand covered F1 (Fig. 13). The Spearman correlations between site sequence identity and covered F1 were −0.258, −0.086, and −0.128 for the overrepresented, underrepresented, and zero-shot groups, respectively. After adjustment, the corresponding correlations were −0.087, 0.115, and 0.095. Thus, binding-site amino-acid identity did not show a stable positive relationship with predictive performance, and the direction of the association changed after adjustment in two groups. These results do not establish that conservation is harmful or rule out memorization of conserved binding sites.

**Figure 13.**
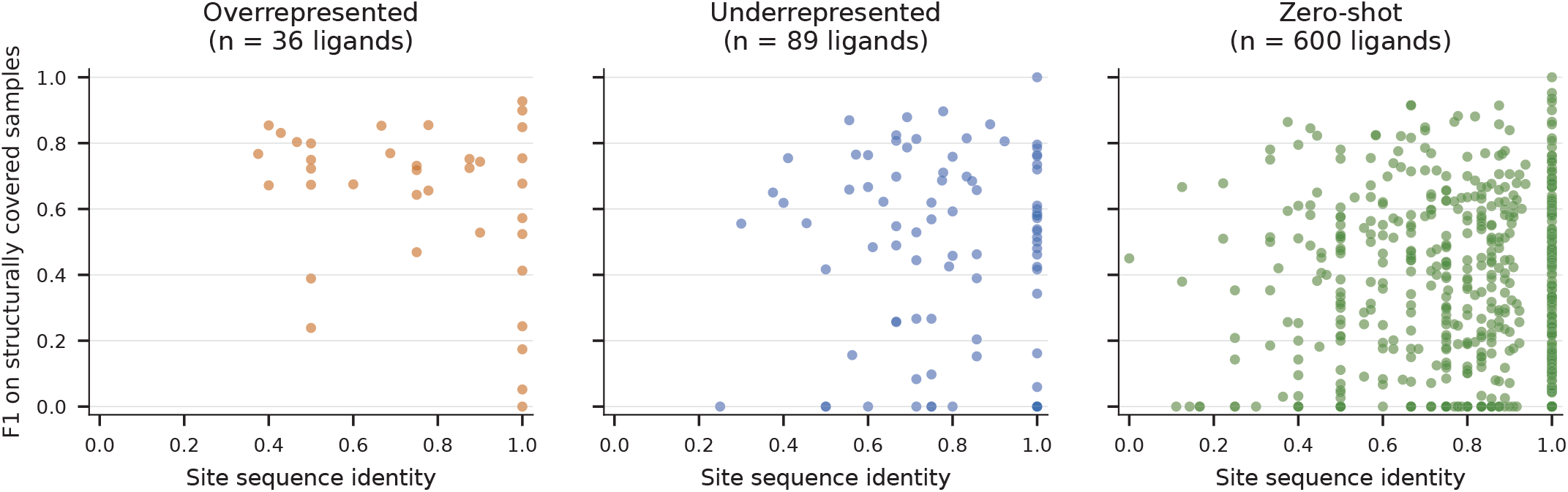
Reference binding-site sequence conservation and performance. Each point represents one of the same 36, 89, or 600 structurally covered ligands. Higher site sequence identity means greater conservation among the accepted reference comparisons. F1 uses only contributing covered samples. These are unadjusted descriptive associations.

## Performance under alternative weighting schemes

We additionally examined whether the summary of Stage 3 performance depended on the weighting scheme (Fig. 14). Using the same saved predictions and evaluation mask, pooled F1, receptor-cluster-balanced F1, and ligand-macro F1 were 0.6579, 0.5338, and 0.5501 for the overrepresented group; 0.4338, 0.3327, and 0.3596 for the under-represented group; and 0.3193, 0.2537, and 0.2379 for the zero-shot group, respectively. For the cluster-balanced metric, residue-level confusion counts were pooled within each globally defined receptor cluster, F1 was calculated per cluster, and represented clusters were given equal weight. Ligand-macro F1 instead gives each ligand equal weight. These metrics therefore provide different weighting summaries of the same Stage 3 predictions rather than results from different models or evidence of a causal performance loss. The lower balanced scores indicate that the weighting scheme affects the reported summary, but do not by themselves establish memorization.

**Figure 14.**
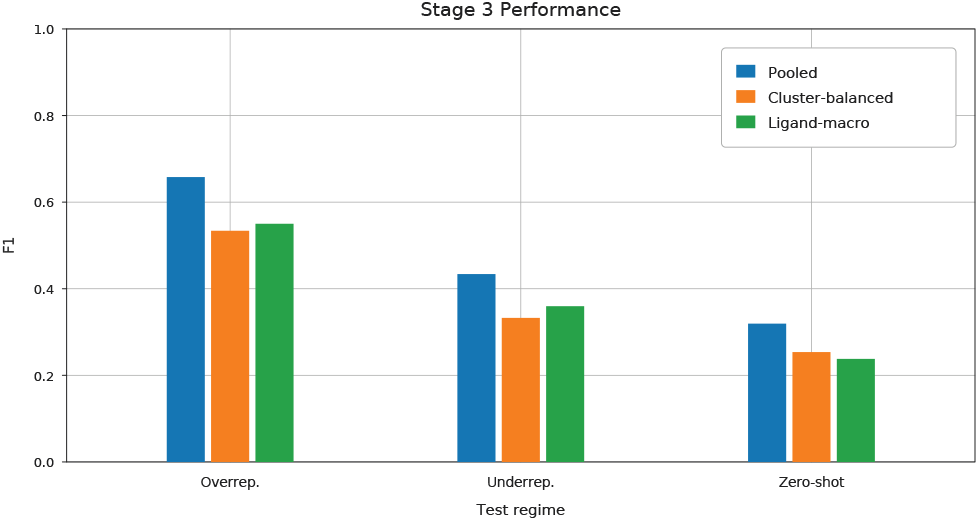
Performance under three weighting rules on the full modified Stage 3 test datasets. Pooled residue-level *F*_1_, receptor-cluster-balanced *F*_1_, and ligand-macro *F*_1_ use the same saved predictions and evaluation mask.

## Matched Linear-Predictor versus Decoder Ablation

To separate the effect of predictor architecture from the effects of multi-ligand training and ligand conditioning, we performed an additional matched ablation on six selected overrepresented ligands: `ZN, ATP, PLP, GLC, FE2`, and `BMA`. For each ligand, we compared the Stage 1 linear predictor with a Stage 2-compatible decoder while retaining ligand-specific training. Each configuration was trained using three paired random seeds, resulting in 36 training runs (6 ligands × 2 predictor architectures × 3 seeds).

### Matched training settings

Both predictor arms used ESM-2 650M, with the final three encoder layers fine-tuned and the earlier encoder layers and embeddings frozen. The shared settings were a batch size of 16, 24 training epochs, a maximum protein length of 768, the same ligand-specific data, paired seeds, initialization, optimizer, learning-rate schedule, validation-based checkpoint selection, and a probability threshold greater than 0.5. The comparison used the canonical modified training, validation, and overrepresented-test files.

The linear arm used the Stage 1 linear predictor. The decoder arm used a Stage 2-compatible four-layer, eight-head decoder with a feedforward width of 2048 and cross-attention to one learned ligand embedding. The ligand embedding was constant across samples within each per-ligand model. Both arms used AdamW with a learning rate of 5 × 10^−5^, *β*_1_ = 0.9, *β*_2_ = 0.98, *ϵ* = 10^−7^, weight decay of 0.01, cosine decay, BF16 computation, and gradient clipping at 1.0.

Both arms reproduced the legacy Stage 1 batch-dependent positive weighting and included padded positions in the loss, checkpoint-selection metrics, and test metrics. This padded-position protocol applies only to the matched six-ligand ablation reported here. The main Stage 1–Stage 3 experiments, including the Stage 1 result in Table 2, excluded padding from the loss, checkpoint-selection metrics, and final evaluation. For the first batch, the positive-class coefficient was set to the negative-to-positive count ratio. For subsequent batches, it was updated as

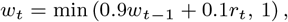

where *r*_*t*_ is the negative-to-positive count ratio in the current batch. This evaluation convention is distinct from the valid-residue Stage 3 diagnostics reported elsewhere.

Under these matched conditions, the decoder achieved a six-ligand ligand-macro *F*_1_ of 0.5567 ± 0.0198, compared with 0.5431 ± 0.0048 for the linear predictor, corresponding to a paired difference of +0.0136 ± 0.0171. Thus, the decoder produced only a modest average improvement in this controlled comparison. This indicates that the larger decoder alone is unlikely to explain the improvement observed in the original Stage 1–Stage 2 comparison. The results are consistent with additional contributions from multi-ligand training and ligand conditioning, although this experiment does not separately quantify those effects.

## Ligand-input control bootstrap analysis

To quantify uncertainty in the ligand-input control experiments, we performed paired bootstrap analysis for each native-versus-control comparison. Differences were calculated as native-SMILES pooled *F*_1_ minus control pooled *F*_1_. Confidence intervals were estimated using 2,000 paired bootstrap resamples of exact receptor-sequence groups within each test set. Native and control observations were resampled together. The resulting differences and 95% confidence intervals are reported in Table 16.

## Architecture Details

**Table 17.** Key hyperparameters for Stage 2 and Stage 3 architectures across different model scales. The decoder activation reported here for the released Stage 3 reference and control evaluations is ReLU.

| **Protein encoder (ESM-2)** | 8M | 35M | 150M | 650M | 3B |
| **Protein encoder dimension** | 320 | 480 | 640 | 1280 | 2560 |
|---|---|---|---|---|---|
| **Decoder dimension** | 320 | 480 | 640 | 1280 | 2560 |
| **Decoder blocks** | 4 |  |  |  |  |
| **Decoder heads** | 8 |  |  |  |  |
| **Decoder FF size** | 2048 |  |  |  |  |
| **Decoder activation (released evaluation)** | ReLU |  |  |  |  |

**Table 18.** Reference Stage 3 configuration. Epoch 14 denotes the best-checkpoint epoch, rather than the total training duration. These settings apply to the reference Stage 3 model.

| **Setting** | **Value** |
|---|---|
| Best-checkpoint epoch | 14 |
| Batch size | 32 samples* |
| Optimizer | AdamW |
| Learning rate | 5 × 10<sup>−5</sup> |
| Scheduler | Cosine annealing |
| Weight decay | 0.01 |
| Maximum protein length | 1,024 residues |
| Maximum SMILES length | 128 tokens, including special tokens |
| Gradient clipping | Maximum norm of 1.0 |
| Protein encoder | Last 3 ESM-2 blocks fine-tuned; remaining blocks and token embeddings frozen |
| Chemical encoder | MoLFormer fully frozen |
| Mixed precision | BF16 |
| Checkpoint selection | Best validation ligand-macro *F*<sub>1</sub> |
| Prediction threshold | Probabilities strictly greater than 0.5 classified as binding |

\*Batch size 32 matches the current single-device configuration and code; its use in the original training still needs confirmation. The current training loop does not use the gradient-accumulation setting.

**Table 19.** Key hyperparameters for Stage 3 architectures across different CLM scales.

| **Chemical encoder** | MolFormer (47M) | UniMol-2 (84M) | UniMol-2 (570M) |
|---|---|---|---|
| **Encoder dimension** | 768 | 768 | 1536 |

**Table 20.** Benchmark of Stage 1 architecture on the ligands of the *overrepresented* test set.

| **Ligand** | *F*<sub>1</sub> |
|---|---|
| ACO | 0.5290 |
| ADP | 0.6351 |
| AGS | 0.5198 |
| AMP | 0.2971 |
| ANP | 0.5012 |
| ATP | 0.5672 |
| BGC | 0.3015 |
| BMA | 0.0000 |
| Ca<sup>2+</sup> | 0.6834 |
| CHL | 0.3651 |
| CLA | 0.5615 |
| CLR | 0.0000 |
| CO | 0.5175 |
| COA | 0.3712 |
| Cu<sup>2+</sup> | 0.7131 |
| FAD | 0.7604 |
| Fe<sup>3+</sup> | 0.6908 |
| Fe<sup>2+</sup> | 0.6226 |
| FES | 0.6481 |
| FMN | 0.7252 |
| FUC | 0.0000 |
| GAL | 0.3359 |
| GDP | 0.7140 |
| GLC | 0.3052 |
| GOL | 0.0219 |
| GSH | 0.6271 |
| GTP | 0.5530 |
| HEC | 0.6713 |
| HEM | 0.6675 |
| MAN | 0.0171 |
| Mg<sup>2+</sup> | 0.5906 |
| Mn<sup>2+</sup> | 0.6044 |
| NAD | 0.6904 |
| NAP | 0.7151 |
| NDP | 0.6444 |
| PLP | 0.7432 |
| PO<sub>4</sub><sup>3−</sup> | 0.3733 |
| SAH | 0.6334 |
| SAM | 0.5323 |
| SF4 | 0.6321 |
| SIA | 0.1011 |
| SO<sub>4</sub><sup>2−</sup> | 0.0000 |
| UDP | 0.4515 |
| Y01 | 0.0000 |
| Zn<sup>2+</sup> | 0.8245 |

**Table 21.** Comparison on the 41 ligands shared between our *overrepresented* test set and the Prot2Token results, out of 45 overrepresented ligands. All values of Prot2Token are reported from the original paper.

| **Ligands** | **Stage 1** | **Stage 2** | **Stage 3** | **Prot2Token** |
|---|---|---|---|---|
| Zn<sup>2+</sup> | 0.8245 | 0.8180 | 0.7747 | 0.7575 |
| Ca<sup>2+</sup> | 0.6834 | 0.6948 | 0.6239 | 0.6474 |
| CLA | 0.5615 | 0.5740 | 0.5252 | 0.4762 |
| FAD | 0.7604 | 0.7833 | 0.7667 | 0.6537 |
| HEM | 0.6675 | 0.6922 | 0.6859 | 0.6796 |
| NAD | 0.6904 | 0.7515 | 0.7118 | 0.6952 |
| ADP | 0.6351 | 0.6930 | 0.6801 | 0.5834 |
| Mg<sup>2+</sup> | 0.5906 | 0.5637 | 0.4985 | 0.4575 |
| NAP | 0.7151 | 0.7696 | 0.7424 | 0.6746 |
| ATP | 0.5672 | 0.6775 | 0.6478 | 0.5050 |
| **Average (top 10)** | **0.6696** | **0.7018** | **0.6657** | **0.6130** |
| HEC | 0.6713 | 0.7525 | 0.7137 | 0.6537 |
| SF4 | 0.6321 | 0.7750 | 0.7086 | 0.5685 |
| FMN | 0.7252 | 0.7966 | 0.8153 | 0.6945 |
| SAH | 0.6334 | 0.7553 | 0.7134 | 0.6503 |
| NDP | 0.6444 | 0.8076 | 0.7859 | 0.6979 |
| ANP | 0.5012 | 0.7078 | 0.6777 | 0.6217 |
| GDP | 0.7140 | 0.8035 | 0.7862 | 0.6465 |
| GLC | 0.3052 | 0.3166 | 0.3058 | 0.2214 |
| PLP | 0.7432 | 0.8311 | 0.8184 | 0.7620 |
| Mn<sup>2+</sup> | 0.6044 | 0.7663 | 0.6876 | 0.7376 |
| **Average (top 20)** | **0.6435** | **0.7165** | **0.6835** | **0.6192** |
| COA | 0.3712 | 0.5134 | 0.4969 | 0.4011 |
| SAM | 0.5323 | 0.7022 | 0.6680 | 0.6252 |
| AMP | 0.2971 | 0.5306 | 0.5011 | 0.4432 |
| BGC | 0.3015 | 0.3514 | 0.3223 | 0.1932 |
| Fe<sup>3+</sup> | 0.6908 | 0.7993 | 0.7551 | 0.6606 |
| MAN | 0.0171 | 0.1538 | 0.0822 | 0.1216 |
| FES | 0.6481 | 0.7693 | 0.7167 | 0.7018 |
| PO<sub>4</sub><sup>3−</sup> | 0.3733 | 0.3970 | 0.3178 | 0.2278 |
| GTP | 0.5530 | 0.6445 | 0.6412 | 0.5461 |
| UDP | 0.4515 | 0.6158 | 0.5994 | 0.5391 |
| **Average (top 30)** | **0.5702** | **0.6602** | **0.6257** | **0.5615** |
| Cu<sup>2+</sup> | 0.7131 | 0.6893 | 0.6328 | 0.5607 |
| GSH | 0.6271 | 0.7759 | 0.7555 | 0.6924 |
| AGS | 0.5198 | 0.6785 | 0.6914 | 0.5301 |
| ACO | 0.5290 | 0.6391 | 0.6034 | 0.5026 |
| GAL | 0.3359 | 0.4616 | 0.4649 | 0.2762 |
| SO<sub>4</sub><sup>2−</sup> | 0.0000 | 0.1807 | 0.2049 | 0.1386 |
| CLR | 0.0000 | 0.0568 | 0.0449 | 0.0373 |
| Y01 | 0.0000 | 0.0981 | 0.0377 | 0.0419 |
| BMA | 0.0000 | 0.2927 | 0.2857 | 0.2273 |
| Fe<sup>2+</sup> | 0.6226 | 0.8965 | 0.8213 | 0.6033 |
| CO | 0.5175 | 0.6510 | 0.7048 | 0.5170 |
| **Average (41 shared ligands)** | **0.5115** | **0.6153** | **0.5858** | **0.5115** |

**Table 22.** Pooled residue-level *F*_1_ values underlying Figure 5, organized by test regime, chemical encoder, and ESM-2 backbone size. “2 unfrozen” and “4 unfrozen” denote fine-tuning of the final two or four chemicalencoder transformer blocks, respectively.

| **Chemical encoder** | **ESM-2 backbone size** |  |  |  |  |
|---|---|---|---|---|---|
|  | **8M** | **35M** | **150M** | **650M** | **3B** |
| **(a) Overrepresented** |  |  |  |  |  |
| MolFormer | 0.5125 | 0.5770 | 0.6153 | 0.6469 | 0.6689 |
| UniMol-2 84M | 0.4962 | 0.5637 | 0.6013 | 0.6241 | 0.6283 |
| UniMol-2 84M<br>(2 unfrozen) | 0.5059 | 0.5632 | 0.5949 | 0.6222 | 0.6492 |
| UniMol-2 84M<br>(4 unfrozen) | 0.5026 | 0.5688 | 0.6104 | 0.6342 | 0.6510 |
| UniMol-2 570M | 0.5039 | 0.5619 | 0.5865 | 0.6268 | 0.6440 |
| **(b) Underrepresented** |  |  |  |  |  |
| MolFormer | 0.2088 | 0.2760 | 0.3200 | 0.3946 | 0.4315 |
| UniMol-2 84M | 0.2019 | 0.2654 | 0.2958 | 0.3278 | 0.3637 |
| UniMol-2 84M<br>(2 unfrozen) | 0.2121 | 0.2481 | 0.2765 | 0.3594 | 0.3716 |
| UniMol-2 84M<br>(4 unfrozen) | 0.2015 | 0.2595 | 0.2890 | 0.3610 | 0.3629 |
| UniMol-2 570M | 0.1991 | 0.2516 | 0.2858 | 0.3348 | 0.3673 |
| **(c) Zero-shot** |  |  |  |  |  |
| MolFormer | 0.1502 | 0.1876 | 0.2402 | 0.3109 | 0.2979 |
| UniMol-2 84M | 0.1444 | 0.2100 | 0.2443 | 0.2530 | 0.2708 |
| UniMol-2 84M<br>(2 unfrozen) | 0.1508 | 0.1904 | 0.2201 | 0.2756 | 0.2881 |
| UniMol-2 84M<br>(4 unfrozen) | 0.1474 | 0.1966 | 0.2311 | 0.2858 | 0.2771 |
| UniMol-2 570M | 0.1417 | 0.1883 | 0.2248 | 0.2594 | 0.2799 |

